# Synergy of color and motion vision for detecting approaching objects in *Drosophila*

**DOI:** 10.1101/2021.11.03.467132

**Authors:** Kit D. Longden, Edward M. Rogers, Aljoscha Nern, Heather Dionne, Michael B. Reiser

## Abstract

Color and motion are used by many species to identify salient moving objects. They are processed largely independently, but color contributes to motion processing in humans, for example, enabling moving colored objects to be detected when their luminance matches the background. Here, we demonstrate an unexpected, additional contribution of color to motion vision in *Drosophila*. We show that behavioral ON-motion responses are more sensitive to UV than for OFF-motion, and we identify cellular pathways connecting UV-sensitive R7 photoreceptors to ON and OFF-motion-sensitive T4 and T5 cells, using neurogenetics and calcium imaging. Remarkably, the synergy of color and motion vision enhances the detection of approaching UV discs, but not green discs with the same chromatic contrast, and we show how this generalizes for visual systems with ON and OFF pathways. Our results provide a computational and circuit basis for how color enhances motion vision to favor the detection of saliently colored objects.

## Main

Color and motion are two visual cues used by many species to identify salient moving objects^1,2^. They are processed largely independently, for example along the ventral and dorsal pathways, respectively, in primates^3,4^. However, color contributes to motion perception in humans and other primates, as indicated by psychophysical experiments using chromatic stimuli lacking luminance contrast^5,6^, allowing the edge motion of objects to be detected even when the background illumination changes to match the luminance of the object. Here, by identifying cellular pathways for color contributing to motion processing in *Drosophila*, we demonstrate how color can additionally boost the motion detection of specifically colored objects, without, surprisingly, improving the motion detection of differently colored objects with the same chromatic contrast.

Like primates, color and motion are initially largely processed along separate pathways in *Drosophila*^7^, and their motion vision was not thought to be influenced by color vision based on studies using blue and green wavelengths^8–11^. However, connectomic studies found that the photoreceptors used for color vision are also connected to cells in the motion pathway^12,13^, and a study using sophisticated genetic manipulations indicated that color inputs expand the spectral range of motion vision through unknown cellular mechanisms^14^. *Drosophila* forages on fruits, flowers, and water droplets that can all have a UV reflectance distinct from background foliage^15,16^, and UV illumination is used commercially to identify ripe or damaged citrus fruit^17^. We therefore explored how color might contribute to motion vision using the behavioral and cellular responses of *Drosophila* to objects defined by UV-green color edges.

*Drosophila* has a visual system well suited to parsing UV and green components of the visual scene. Under every eye facet (ommatidium) are eight photoreceptors (Fig. 1a). The outer six, R1-6, are sensitive to a broad range of wavelengths, especially UV and green, and provide the luminance signal for pathways processing motion vision^18^. The inner two, R7 and R8, provide additional wavelength sensitivity for color vision^19–22^. The R7 and R8 neurons are paired and come in two flavors, determined by the rhodopsins expressed, across most of the eye: one sensitive to short wavelength UV and blue respectively (in so-called pale ommatidia), and the other to long wavelength UV and green (in so-called yellow ommatidia). R1-6 and R7-8 project to different visual neuropils, the lamina and medulla, respectively^7,23,24^, and like vertebrates, flies have visual processing pathways for the movement of bright edges relative to the background (ON-motion), and dark edges relative to the background (OFF-motion). The T4 and T5 cell types are directionally selective for ON- and OFF-motion, respectively^25^, and the circuitry connecting the photoreceptors to T4 and T5 neurons has been described in considerable detail^12,26,27^.

**Figure 1.**
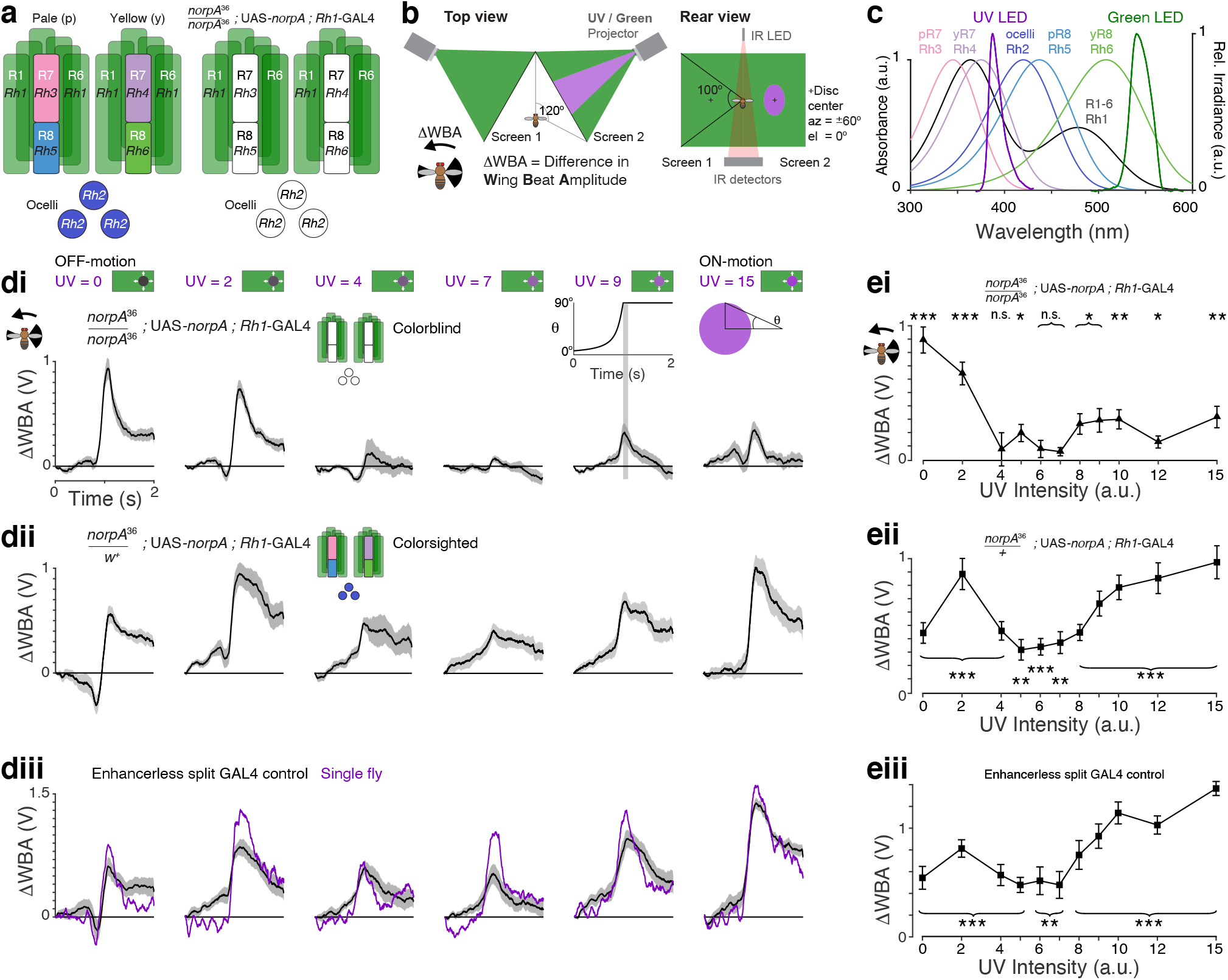
Color contributes to motion vision in *Drosophila*’s responses to expanding discs. **a**. Rhodopsin expression in pale and yellow ommatidia and ocelli in wild type flies (left) and in colorblind *norpA*^36^ Rh1-rescue flies in which the wildtype *norpA* has been rescued only in the outer photoreceptors (right); non-functional photoreceptors are indicated by white. **b**. The behavioral setup, as seen from above (left) and behind (right). Two customized LED projectors displayed UV and green stimuli onto Teflon screens, covering a view of 240^°^ azimuth and 100^°^ elevation. Turns were measured as the difference in the Wing Beat Amplitude, ΔWBA, between the left and right wings, using infrared (IR) illumination from above and IR detectors below. Approaching discs were centered at ±60^°^ azimuth, 0^°^ elevation. **c**. Spectral sensitivity (absorbance) of rhodopsins^19^, and irradiance spectra of the UV and green projector LEDs. **d**. Responses to expanding UV discs of different genotypes (di-iii) for selected UV intensities given in purple and illustrated above each panel column. Stimulus time course is shown above the top row panel for UV = 9. The radial angle of the disc seen by the fly, θ, increased over 1 s and was parameterized by the ratio of the radius to the velocity, r/v = 120 ms, which is approximately the rate of an approaching human hand trying to swat a fly. The fully expanded disc remained for 1 s before the screen returned to green. For all rows, *N* =10 flies, and mean ±SEM shown. **di**. Colorblind *norpA*^36^ Rh1-rescue flies with *norpA* expression rescued in R1-6. **dii**. Colorsighted *norpA*^36^ Rh1-rescue control flies with heterozygous expression of *norpA* in R1-8 and ocelli. **diii**. Enhancerless split GAL4 > wild type *DL*. The mean responses for a single fly are shown in purple. **e**. Responses for each UV intensity presented, for flies shown in (**d**). Responses are measured as the mean response in the 100 ms after the disc has fully expanded, as indicated by the gray stripe in the stimulus diagram inset (above UV = 9 column in **d**). For all rows, *N* =10 flies, and mean ±SEM shown. Student’s *t*-test was used to identify responses significantly different from zero, with FDR correction for 11 comparisons. Asterisks indicate significance level: *\* p* < 0.05, *\*\* p* < 0.01, \*\**\* p* < 0.001, n.s. not significant. Genotypes for all flies used in behavioral experiments are listed in Table 1.

To test if color contributes to motion vision in flies, we developed a custom projector to display spatially precise UV and green patterns that corrected for impedimentary short-wave scattering, and then measured behavioral responses to expanding discs that varied between dark and bright UV with a green background. Flies responded to all intensities of UV discs, indicating that color contributed to motion vision. Remarkably, responses to ON-motion were much more sensitive to UV than those to OFF-motion in a range of *Drosophila* species. To identify the cells responsible, we developed genetic reagents to manipulate the function of classes of photoreceptors, and to identify cells downstream of R7, we used 2-photon calcium imaging of neuronal activity. Our analysis of these results generated the counterintuitive prediction that the contribution of color to motion vision would not support the detection of green discs seen against UV, even though both UV and green discs have the same chromatic contrast, and remarkably, this was the case. Finally, we determined how the mechanism can be employed in other visual systems to detect objects of other colors. These findings identify cellular pathways linking color photoreceptors and motion-sensitive cells underlying the synergy of color to motion vision, and show how the mechanism can be selective for UV objects in *Drosophila*, and be employed for color-selectivity in other visual systems.

## Results

### Color contributes to motion vision in *Drosophila*’s responses to expanding discs

Testing for a contribution of color to motion vision requires a spatially precise display system. We customized a projector to display UV-green patterns, with spectra matched to the spectral tuning of the photoreceptors (Fig. 1b-c), and matched the green irradiance to the UV using a luminance mask (Extended Data Fig. 1a-d). Without this calibration, scattering in the projection screen varies the ratio of green to UV light by a factor of up to 5 (Extended Data Fig. 1a-e). As a consequence, the UV intensity is measured in levels, ranging 0-15, rather than the absolute irradiance, and green levels were fixed for all stimuli (see Methods). We presented flying, tethered flies with UV discs expanding out of a fixed green background, as if approaching at a constant velocity from one side. Expanding discs are an ideal stimulus to probe for a contribution of color to motion vision because they present just one kind of color edge to be processed by the ON and OFF-motion pathways: a dark UV disc presents only the OFF-motion of dark UV moving into green, while a bright UV disc presents only the ON-motion of bright UV moving into green. The flies readily turn away from the approaching discs, and the behavior is reliably measured by optically recording the difference in the amplitude of the left- and right-wing beats (Fig. 1b, ΔWBA).

We used colorblind flies (Fig. 1a, right) to establish when UV and green were matched for the luminance channel that is driven by the R1-6 (outer) photoreceptors. In *norpA*^36^ flies, the phototransduction pathway is not functional for motion vision, and we rescued the function of R1-6 by expressing the wild-type *norpA* in the Rh1-expressing cells using the GAL4-UAS system. These flies are colorblind because phototransduction is only functional for photoreceptors expressing one light-sensitive rhodopsin, Rh1 (Fig. 1a). No matter what crosstalk exists between downstream pathways, all visual inputs are constrained by one spectral tuning in these flies.

When presented with an approaching UV disc, the colorblind flies robustly turned away when it was dark (Fig. 1di, UV = 0), or bright (Fig. 1di, UV = 15). At intermediate UV intensities, their peak response reduced to near zero (Fig. 1di, UV = 7), and the flies behaved as though they did not see the motion of the disc, indicating the disc and the background were effectively isoluminant (Fig. 1ei; UV = 4, 6 − 7; p > 0.05, *t*-test with false discovery rate (FDR) correction for multiple comparisons, *N* = 10). When the activity of all the photoreceptors was restored in heterozygous control (colorsighted) flies, they turned away from all approaching UV discs, including the intermediate intensities that were isoluminant with the green background for the colorblind flies (Fig. 1dii, eii; UV = 4 − 7; *p* < 0.01, *t*-test with FDR correction, *N* = 10). The control flies turned to avoid approaching discs for all UV intensities and lacked a null response to isoluminant discs, demonstrating that color contributed to motion vision.

Genetic control flies prominently used in our further experiments also demonstrated a contribution of color to motion vision by responding to isoluminant UV discs (Fig. 1diii, eiii; Enhancerless split GAL4 > wild type *DL*, *p* < 0.01, *t*-test with FDR correction, *N* = 10), and also responded to discs with an intermediate intensity (UV = 6) over a wide range of approaching speeds (Extended Data Fig. 1f). Avoidance of expanding visual patterns requires functional T4 and T5 cells, the ON- and OFF-motion directionally selective cells^28^, and these cells were also required for the contribution of color to motion vision: when they were silenced by expressing the inwardly rectifying potassium channel Kir_2.1_ ^29^, responses to expanding UV discs were abolished (Extended Data Fig. 1g, h; responses compared to zero, *t*-test: *p* > 0.0.5, *t*-test with FDR correction, *N* = 10).

### Behavioral responses to ON- and OFF-motion differ in their sensitivity to UV and green

As motion processing in flies is divided into pathways for ON- and OFF-motion, we tested whether the processing of ON- and OFF-motion may be differently sensitive to UV and green by measuring turning responses to competing moving edges. We divided the fly’s visual panorama into eight windows along the horizon, and in each window presented the same stimulus, illustrated in Fig. 2a. For competing ON-motion, UV and green patches expanded horizontally in opposing directions at the same time. Flies turn to follow movement of the visual scene and they follow the edges with the greatest contrast. They fly straight, on average, when the contrast of the edges balance: this is the isoluminance level. For competing OFF-motion, the UV and green patches contracted horizontally within each window − the identical stimulus frames were presented, just in the reverse frame sequence as for the ON-motion.

**Figure 2.**
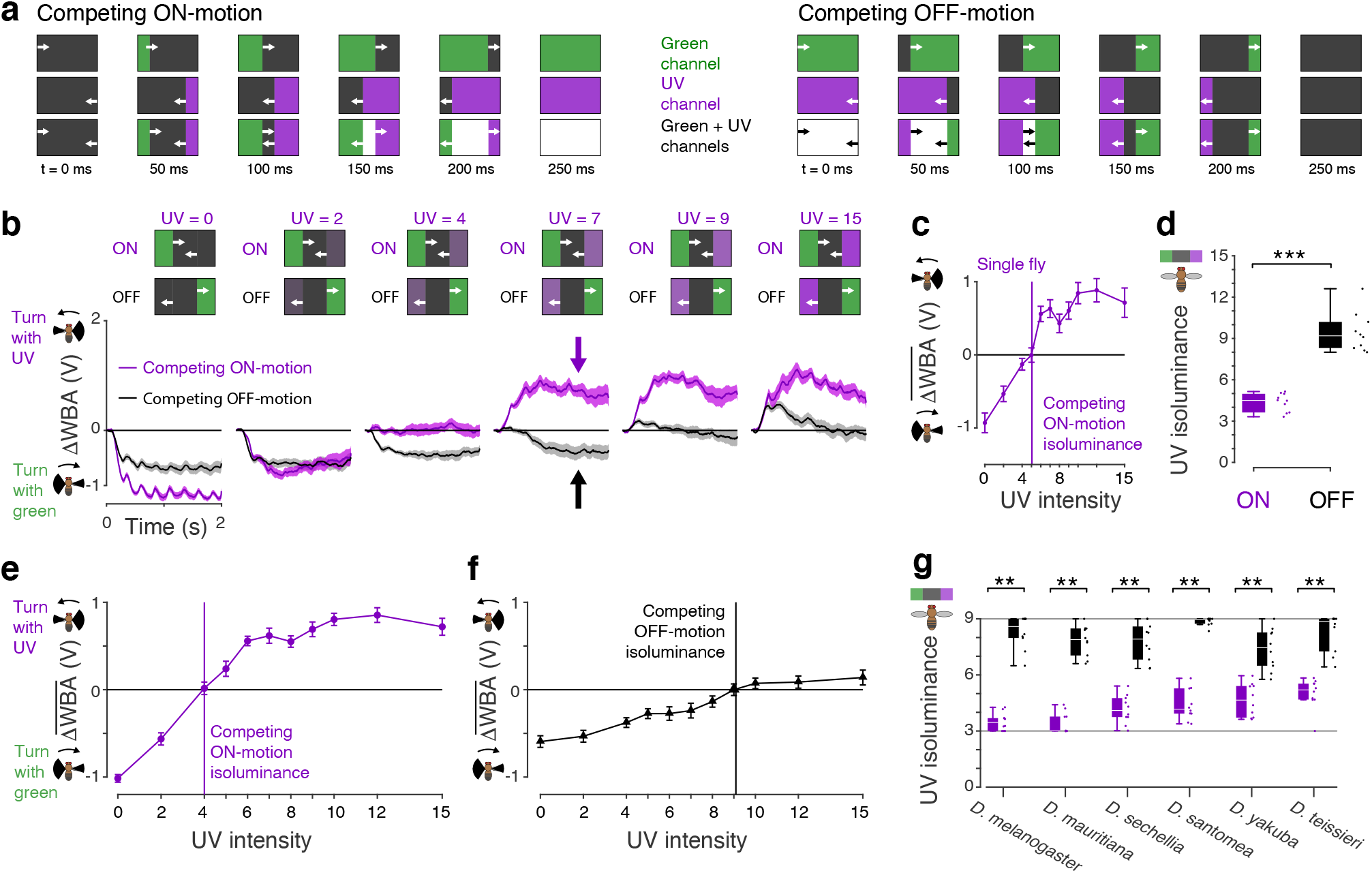
Behavioral responses to ON- and OFF-motion differ in their sensitivity to UV. **a**. Diagrams of competing motion stimuli. The display was divided into 8 windows of 30° azimuth each, and diagrams illustrate sample frames of the stimuli within each window for the times indicated at the bottom. Top row shows the green channel component, middle row the UV component, and the bottom row the summation of the two that were the stimuli presented to the flies. Each stimulus cycle lasted 250 ms (4Hz) and were shown for 2 s. **b**. Turning responses of flies our primary data genotype, enhancerless split GAL4 crossed with type (DL), to competing ON- (purple) and OFF-motion (black). Time traces are shown for selected UV intensities indicated in purple above every panel. The panel for UV = 7 is highlighted: during the ON-motion stimuli, the flies turn to follow the moving UV edges (purple arrow), indicating the UV edges have greater ON-motion contrast than the green edges; during the OFF-motion stimuli, the flies turn to follow the motion of green edges (black arrow), indicating the UV edges have less OFF-motion contrast than the green edges. *N*_ON_ =10, *N*_OFF_ =10 (mean ± SEM). Data from the same flies are plotted in panels (d-f). **c**. Example isoluminance calculation, for a single fly’s response to competing ON-motion. The mean ± SEM response over the 2 s of the stimulus, and 5 trials, is shown. The isoluminance level is the lowest UV intensity when the response is greater than zero, using linear interpolation between stimulus intensities. **d**. Isoluminance levels of populations of flies for competing ON- and OFF-motion, compared using a two-sample student’s *t*-test, *** indicates *p* < 0.001, *N*_ON_ =10, *N*_OFF_ =10. Boxplots indicate the median and quartile ranges, and whiskers indicate the extent of data points within an additional 1.5 × quartile range, conventions used for the boxplots in all the figures. **e**. Mean response of all flies to competing ON-motion. The isoluminance level of the mean response is indicated by the vertical line. *N*_ON_ =10 (mean ± SEM). **f**. Mean response of all flies to competing OFF-motion. The isoluminance level of the mean response is indicated by the vertical line. *N*_OFF_ =10 (mean ± SEM). **g**. Isoluminance levels of different *Drosophil*a species, for competing ON- (purple) and OFF-motion (black), using a compact protocol where both isoluminance levels were measured in the same flies. The luminance was restricted to the range 3−9, the gray lines at UV = 3 and UV = 9 indicate these bounds. For all species, Wilcoxon signed-rank test comparing the isoluminance levels for competing ON- and OFF-motion, ** indicates *p* < 0.01, *N* = 10 flies. Boxplot conventions are as in panel (d). Genotypes of all flies used in behavioral experiments are listed in Table 1.

When shown dark UV edges, genetic control flies turned with the green edges (Fig. 2b; UV = 0), and they turned with the UV edges when they were bright (Fig. 2b; UV = 15). The UV intensity when they switched from turning with green to turning with UV was different for ON- and OFF-motion: it was ∼4 for ON-motion (Fig. 2b; purple trace, UV = 4), and ∼9 for OFF-motion (Fig. 2b; gray trace, UV = 9). For UV intensities between these values, flies responded to the same intensity levels differently, depending on whether they were viewing ON- or OFF-motion. For example, the flies followed the UV edge for a UV intensity of 7 during ON-motion (Fig. 2b, purple arrow), while for the same intensity they followed the green edge for OFF-motion (Fig. 2b, gray arrow). Therefore, although the intensities of the moving UV and moving green edges are the same, they can drive different responses in the ON- and OFF-motion pathways, depending on the stimulus time-history.

Individual flies had identifiable isoluminance levels where ΔWBA = 0 (Fig. 2c). We compared the isoluminance levels for ON- and OFF-motion, measured in separate groups of flies, and they were significantly different (Fig. 2d; two-sample student’s *t*-test, *p* < 0.001, *N*_ON_ = 10, *N*_OFF_ = 10). Responses to ON-motion were twice as sensitive to UV as to OFF-motion, with a median isoluminance level for ON-motion of 4.5, and 9.2 for OFF-motion (Fig. 2d). We also calculated the isoluminance level of the average, population response to ON-motion, which was 4.0 (Fig. 2e), and of the average, population response to OFF-motion, which was 9.1 (Fig. 2f).

The UV-sensitivity for ON-motion did not depend on the sex of the fly and replicated across setups (Extended Data Fig. 2a). As was the case for the expanding discs, it did require functional T4 and T5 cells because when these cells were silenced by expressing *Kir*_2.1_, the responses to ON- and OFF-edges were largely abolished (Extended Data Fig. 2b, c). We measured the ON- and OFF-motion isoluminance levels in five more *Drosophila* species, and in all the species ON-motion was more sensitive to UV than for OFF-motion (Fig. 2g; for all species *p* < 0.01, Wilcoxon signed-rank test, *N*_Flies_ = 10). These results show that the difference in ON- and OFF-motion UV-sensitivity were not particular to *Drosophila melanogaster* and are shared between other *Drosophila* species.

### R7 photoreceptors support ON-motion UV-sensitivity

To identify which photoreceptors are responsible for the differences between competing ON- and OFF-motion UV-sensitivity, we rescued different combinations of photoreceptors in blind *norpA*^36^ mutant flies. The R1-6 photoreceptors are required because they support the luminance processing in the lamina^18^, and we additionally rescued pale or yellow R7s or R8s, individually or in combination (Fig. 3a, Extended Data 3a). For all the genetic controls for these experiments, there was a significant difference between the isoluminance levels for competing ON-motion (*I*_ON_) and for competing OFF-motion (*I*_OFF_) (Fig. 3a, gray boxplots and data points; Wilcoxon signed-rank test, *p* < 0.01, *N* = 10; the individual values of *I*_ON_ and *I*_OFF_ are shown in Extended Data Fig. 3a).

**Figure 3.**
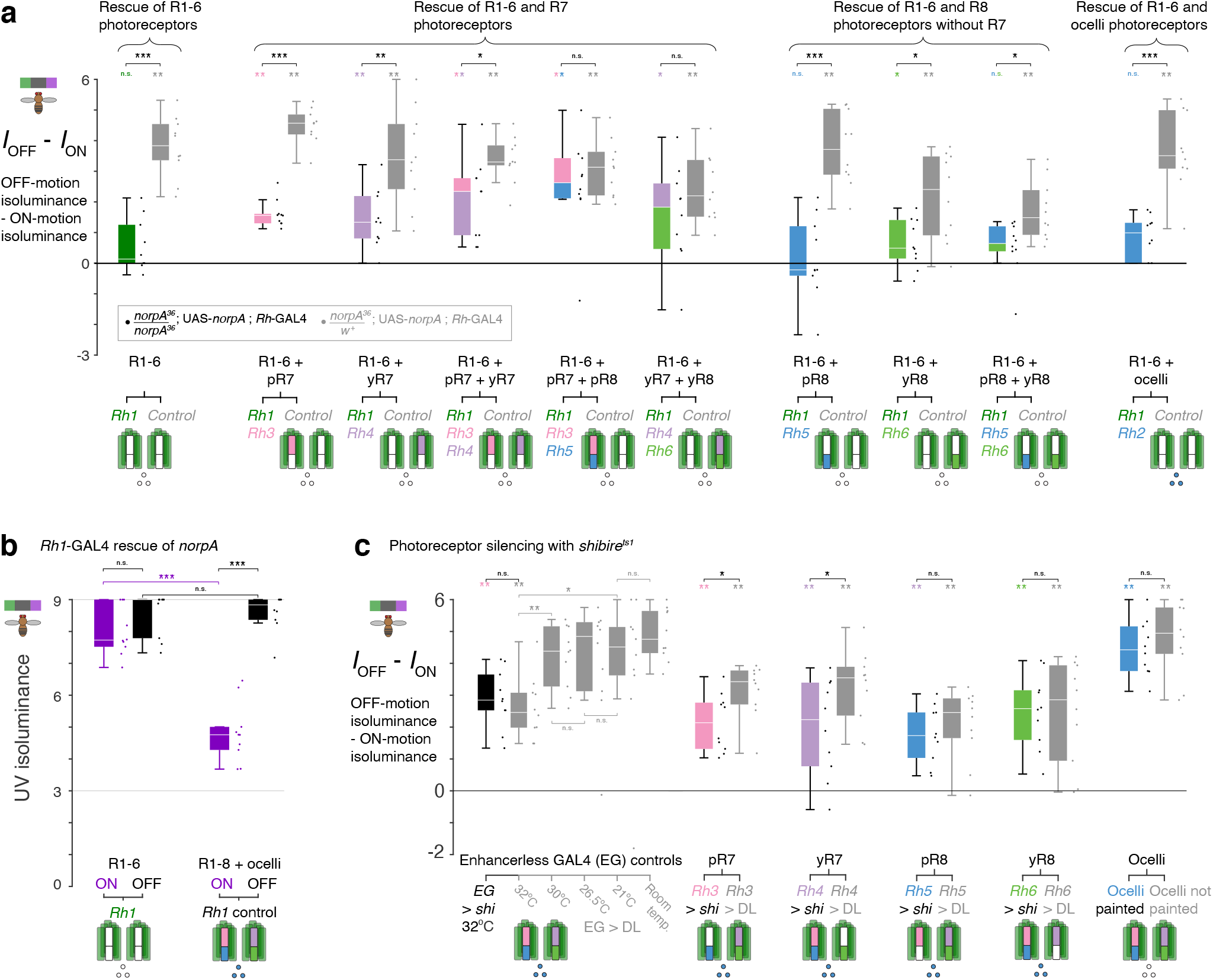
R7 photoreceptors support ON-motion UV-sensitivity. **a**. Differences between the isoluminance levels for competing OFF-motion (*I*_OFF_) and ON-motion (*I*_ON_) of homozygous *norpA*^36^ flies with the function of different combinations of photoreceptors rescued using rhodopsin-GAL4 driven expression of UAS-*norpA* (colored boxplots), and genetic controls (gray boxplots), using the compact protocol where both isoluminance levels were measured in the same flies. Below each pair of boxplots, for every rescue genotype and control, a diagram visually indicates the photoreceptors rescued (colored) and not rescued (white). Above each individual boxplots are statistical tests of whether *I*_OFF_ and *I*_ON_ come from the same distribution, using a paired Wilcoxon signed rank test, *N* = 10 for all genotypes. For comparing between rescue genotypes and controls, we used two sample Wilcoxon rank sum tests. Asterisks indicate significance level: *\* p* < 0.05, *\*\* p* < 0.01, \*\**\* p* < 0.001, n.s. not significant. Boxplot conventions are as in Fig. 2d. The values of *I*_OFF_ and *I*_ON_ for all genotypes of rescued R7 and R8 cells are plotted in Extended Data Fig. 3a. **b**. The values of *I*_OFF_ (black) and *I*_ON_ (purple) for *Rh1*-GAL4 rescue of *norp*A in R1-6 photoreceptors; the pairwise comparisons between *I*_OFF_ and *I*_ON_ contained in this data are shown in panel (a). We used paired Wilcoxon signed rank test to compare *I*_OFF_ and *I*_ON_ within genotypes, and two sample Wilcoxon rank sum tests to compare *I*_OFF_ or *I*_ON_ between rescue and control genotypes, with *N* = 10. Plotting conventions for asterisks as in panel (a). **c**. Differences between the isoluminance levels for competing OFF-and ON-motion (*I*_OFF_ -*I*_ON_), for flies with specific photoreceptor classes silenced using rhodopsin-GAL4 driven expression of UAS-*shibire*^ts1^ (colored boxplots), and controls (gray boxplots), heated to 32°C. The first boxplots show genetic controls that demonstrate the effect of temperature: enhancerless GAL4 crossed with *shibire* heated to 32°C (black), and enhancerless GAL4 crossed with wild type *DL* heated to different temperatures, and not at all (‘room temperature’). For the ocelli, we painted the ocelli with black paint and used enhancerless GAL4 crossed with wild type DL flies. We used paired Wilcoxon signed rank test to compare *I*_OFF_ and *I*_ON_ within genotypes, and two sample Wilcoxon rank sum tests to compare *I*_OFF_ or *I*_ON_ between rescue and control genotypes, with *N* = 10. Plotting conventions for asterisks as in panel (a). The values of *I*_OFF_ and *I*_ON_ for all genotypes of silenced and control flies are plotted in Extended Data Fig. 3c. Genotypes for all flies used in behavioral experiments are listed in Table 1.

When we rescued only R1-6, the difference in the isoluminance levels for competing ON- and OFF-motion (*I*_OFF_ - *I*_ON_) was abolished (Fig. 3a; Wilcoxon signed-rank test R1-6, *p* = 0.11, *N* = 10). Rescuing any R7 cell in addition to R1-6 restored the difference (Fig. 3a; Wilcoxon signed-rank test *p* < 0.05, *N* = 10). The largest effects were for the rescue of R1-6 and both R7s (Fig. 3a; Wilcoxon signed-rank test R1- 6 + pR7 + yR7 *p* = 0.002, median *I*_OFF_ - *I*_ON_ = 2.3, *N* = 10), and for the rescue of R1-6 and R7 coupled with its pale or yellow R8 partner (Fig. 3a; Wilcoxon signed-rank test R1-6 + pR7 + pR8 *p* = 0.004, median *I*_OFF_ - *I*_ON_ = 2.6, *N* = 10; R1-6 + yR7 + yR8 *p* = 0.01, median *I*_OFF_ - *I*_ON_ = 1.8, *N* = 10).

When we rescued R1-6 and R8 cells alone, without their pale or yellow R7 partner cells, there was only a small or no effect on *I*_OFF_ - *I*_ON_ (Fig. 3a). Rescuing R1-6 and pale R8 cells had no significant effect (Fig. 3a; Wilcoxon signed-rank test R1-6 + pR8 *p* > 0.05, *N* = 10), and rescuing R1-6 and yellow R8 resulted in only a small difference (Fig. 3a; Wilcoxon signed-rank test, R1-6 + yR8 *p* = 0.049, median *I*_OFF_ - *I*_ON_ = 0.5, *N* = 10) that was not maintained when we additionally rescued pR8 cells (Fig. 3a; Wilcoxon signed-rank test R1-6 + pR8 + yR8 *p* > 0.05, *N* = 10). Likewise, rescuing R1-6 and the ocelli photoreceptors had no significant effect (Fig. 3a; Wilcoxon signed-rank test R1-6 + ocelli p > 0.05; *N* = 10).

These results indicate that the R7 cells contribute to the difference in the isoluminance levels for competing ON- and OFF-motion. When only R1-6 were rescued the isoluminance level for competing OFF-motion was not affected, compared to controls (Fig. 3b; Wilcoxon rank sum test *p* > 0.05, *N* = 10), while the isoluminance level for ON-motion was significantly increased, compared to controls (Fig. 3b; Wilcoxon rank sum test *p* < 0.001, *N* = 10). Therefore, the difference in the isoluminance levels for competing ON- and OFF-motion is mainly from R7-8 affecting the UV-sensitivity of responses to competing ON-motion. Together these rescue experiments indicate that R7 photoreceptors augment the behaviorally measured ON-motion sensitivity to UV (Fig. 2).

To further support the cellular basis for these findings, we silenced photoreceptors by expressing *shibire^ts^*^1^, a temperature-sensitive mutation of the gene encoding dynamin that inhibits synaptic transmission by blocking vesicle endocytosis^30^ (Fig. 3c, Extended Data Fig. 3b-c). When *shibire* is expressed in R1-6 cells and the flies are tested at an elevated temperature, they are motion blind, and display no directional tuning to the competing motion stimuli (Extended Data Fig. 3b; R1-6 ON/OFF *Rh1* > *shi* T = 32°C, paired *t*-test *p* > 0.05, *N* = 10).

When we silenced either pale or yellow R7 photoreceptors, the difference between the isoluminance levels for competing ON- and OFF-motion was reduced, compared to genetic controls also heated to 32 °C (Fig. 3c; Wilcoxon rank sum test, pR7 *p* = 0.02, yR7 *p* = 0.045, *N* = 10). Silencing R8 photoreceptors had no significant effect on *I*_OFF_ - *I*_ON_, compared to controls (Fig. 3c; Wilcoxon rank sum test, pR8 *p* = 0.38, yR8 *p* = 0.97, N = 10). To silence ocelli photoreceptors, we painted the ocelli of genetic control flies with black paint, and this also had no effect, compared to flies with unpainted ocelli (Fig. 3c; Wilcoxon rank sum test, Ocelli *p* = 0.47, *N* = 10).

Surprisingly, we noted that heating control flies selectively affected the isoluminance level for ON-motion (Extended Data Fig. 3c; comparisons between T = 21°C and T = 32°C for *Rh1* > *DL* flies, Wilcoxon rank sum test *p* = 0.001, *N* = 10), but not the isoluminance level for OFF-motion (Extended Data Fig. 3b; comparisons between T = 21°C and T = 32°C for *Rh1* > *DL* flies, Wilcoxon rank sum test *p* = 0.27, *N* = 10). As a result, we quantified how increasing the heat affects the difference in the isoluminance levels for competing ON- and OFF-motion in genetic control flies (Fig. 3c; Enhancerless GAL4 (EG) > *DL*), and verified that expression of *shibire* had no additional effect when compared to our genetic control flies at 32 °C (Fig. 3c; comparison between EG > *DL* and EG > *shi*, Wilcoxon rank sum test *p* = 0.52, *N* = 10). We therefore used genetic control flies at 32 °C for comparisons (gray boxplots in Fig. 3c).

Taken together, these results clearly show that R7 cells play a pivotal role in supporting the different spectral sensitivities of behavioral responses to ON- and OFF-motion: rescuing the function of R7 cells enabled behavioral responses to ON-motion to be more sensitive to UV, as compared for OFF-motion (Fig. 3a, b), and silencing R7 cells reduced this difference (Fig. 3c).

### The ON- and OFF-motion directionally selective T4 and T5 cells differ in their sensitivity to UV

The T4 and T5 cell types, that are directionally selective for ON- and OFF-motion, respectively, were required for both the flies’ turns away from expanding UV discs on a green background (Extended Data Fig. 1g-h), and the difference in their UV-green isoluminance levels for competing ON- and OFF-motion (Extended Data Fig. 2b-c). Our expectation, based on prior behavioral studies in *Drosophila*, was that T4 and T5 should have identical wavelength sensitivity^10,11,14^, but based on our behavioral results implicating ON and OFF pathway differences, we thought it was essential to evaluate this prediction by measuring the principal motion sensing neurons in each pathway. We therefore examined the responses of T4 and T5 cells to UV discs expanding out of a green background (Fig. 4).

**Figure 4.**
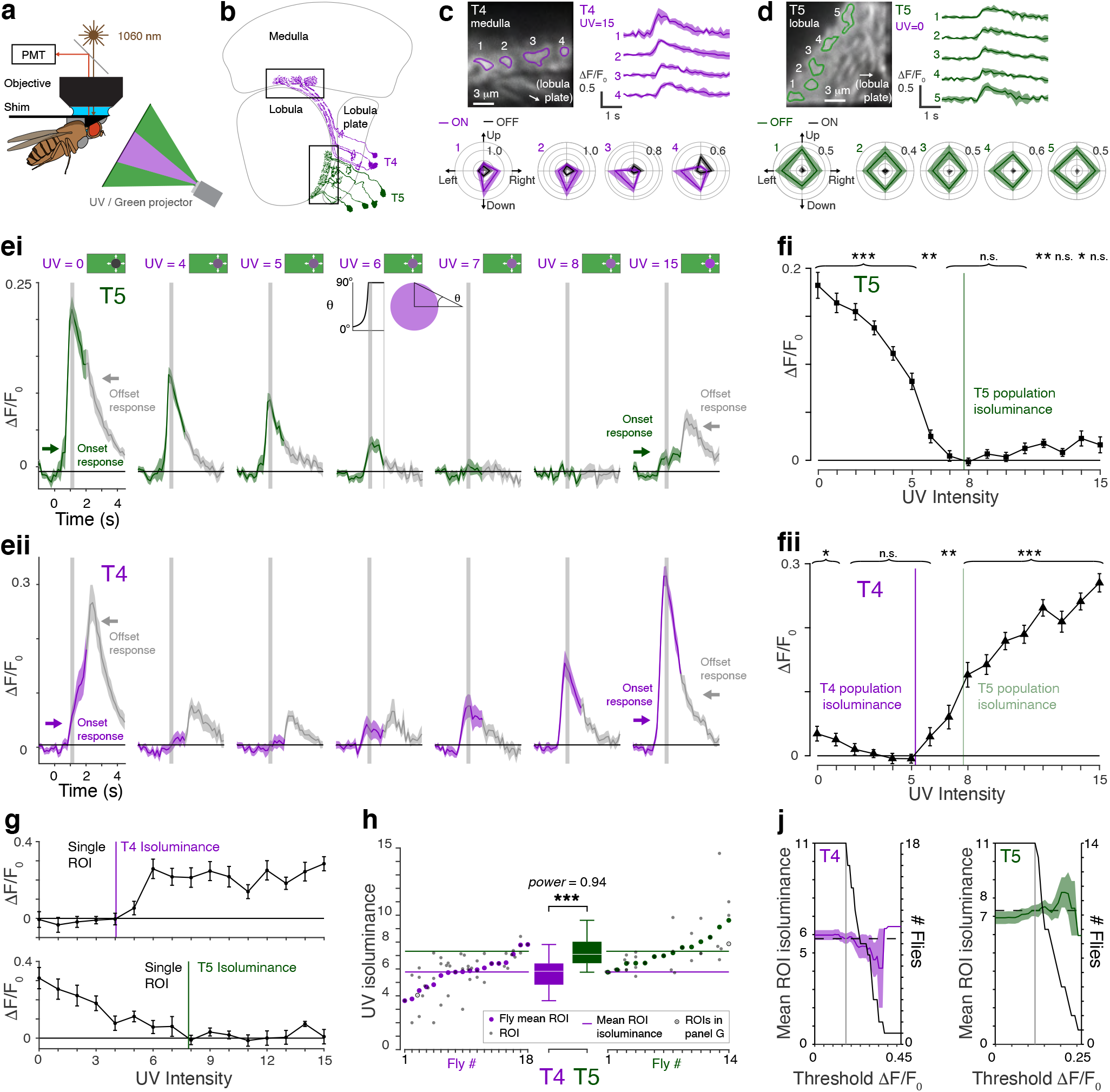
The ON- and OFF-motion directionally selective T4 and T5 cells differ in their sensitivity to UV. **a**. Diagram of imaging setup. **b**. Anatomical diagram of T4 (purple) and T5 cells (green) adapted from^7^. T4 cells were imaged in the medulla and T5 cells in the lobula, indicated by schematic black boxes. **c**. Example recording of T4 cells. Image (top left) shows mean fluorescence for one fly over 5 trials for expanding UV discs, UV = 15. Identified ROIs of columnar units are outlined (purple) and numbered, with corresponding time traces of mean ± S.D. ΔF/F_0_ responses to the stimuli to the right. Responses to unidirectional green edge ON (purple) and OFF-motion (black) stimuli are shown in polar plots below. Some but not all ROIs are directional and so ROIs likely represent multiple cells. **d**. Example recording of T5 cells, organized as in (c), except UV = 0 in the top panels, and responses to directional green edge OFF (green) and ON-motion (black) stimuli in polar plots below. As for T4 cells, ROIs likely represent multiple cells. **e**. Calcium activity responses of T5 (ei) and T4 (eii) ROIs to UV discs expanding from a green background (mean ± SEM), for selected intensities of UV, indicated above the panels. Stimulus time course shown above panel for UV = 6. Traces are colored during the stimulus (T5 green, T4 purple) and switch to gray when the screen returns to background green so that stimulus offset responses can be identified. The mean responses in panel (f) are measured in the 150 ms (two frames) after expansion of the disc, indicated by vertical gray stripe. ROIs averaged for each fly: T4 *N*_Flies_ = 18, *N_ROI_* = 46; T5 *N*_Flies_ = 14, *N_ROI_* = 34. **f**. Mean responses of T5 ROIs (fi) and T4 ROIs (fii) measured in the 150 ms after the disc has expanded, mean ± SEM across flies shown. Colored lines indicate population isoluminance of T4 (fii, purple) and T5 (fi green, and for comparison fii pale green) ROIs. Student’s *t*-test was used to identify responses significantly different from zero, with FDR correction for 16 comparisons. Asterisks indicate significance level: *\* p* < 0.05, *\*\* p* < 0.01, \*\**\* p* < 0.001, n.s. not significant. **g**. Calculation of isoluminance level for single example T4 (top) and T5 (bottom) ROIs. **h**. Mean isoluminance levels of ROIs (gray dots) and flies (color dots); example ROIs in panel G are plotted as a gray dot inside a circle. Colored horizontal lines indicate mean ROI isoluminance levels. The isoluminance levels of T4 ROIs were significantly lower than those of T5 ROIs (*p* < 0.001, two-sample *t*-test, *N*_T4, flies_ = 18, *N*_T5, flies_ = 14; *power* = 0.94), Asterisks indicate significance level: \*\**\* p* < 0.001. Boxplot conventions are as in Fig. 2d. **j**. ROIs were identified using semi-automatic procedure (see Methods), and the threshold used to identify unresponsive ROIs for each cell type are indicated by the vertical gray line. Horizontal dashed line shows mean isoluminance calculated with this threshold. Colored lines and shading indicate mean ROI isoluminance level, averaged over flies ±SEM, as the exclusion threshold is varied. Black lines indicate number of flies with ROIs above the threshold. As the threshold increases, the difference in the T4 and T5 isoluminances remains stable or increases, while statistical power decreases. Genotypes for all flies used in imaging experiments are in Table 2.

We used two photon imaging of intracellular calcium to monitor the activity of the cells. To avoid overlap in the spectral sensitivity of the calcium indicator with the UV and green display, we used the red genetically encoded indicator jRGECO1a ^31^ (Fig. 4a), and added a short-pass wavelength filter, blocking wavelengths longer than green, to a replica of the projector setup used for the behavioral experiments. We imaged T4 dendrites in the medulla and T5 dendrites in the lobula, locations where the cells can be unambiguously identified from a shared driver line (Fig. 4b; Table 2 lists all genotypes for the stocks used in the imaging experiments) and used expanding discs that expanded to a radius of 30° to identify responsive regions of interest (ROIs). As expected^25,32^, T4 ROIs responded preferentially to ON-motion edges (Fig. 4c), and T5 ROIs to OFF-motion (Fig. 4d). Our analysis of T4 and T5 responses is focused on the time window immediately following full disc expansion (Fig. 4eii; gray vertical stripe from *t* = 1 to 1.15 s), corresponding to when behavioral responses peaked (Fig. 1di).

The T5 ROIs responded strongly to black, unilluminated UV discs (Fig. 4ei, UV = 0), consistent with these discs being defined by expanding OFF-edges. For brighter UV discs, the T5 responses declined until there was no significant response for an intensity of UV = 7 (Fig. 4ei). Although T5 cells responded strongly to OFF-motion, they also had small responses to high contrast ON-motion^25,33–35^ and in addition they respond to the OFF-like cessation of ON-motion (Fig. 4ei, UV = 15, offset response indicated by color change to gray). The T4 responses to bright UV discs were strong (Fig. 4eii, UV = 15), consistent with these discs containing expanding ON-edges. For dimmer UV discs, the T4 calcium activity responses were weaker, until there was no significant response for an intensity of UV = 5 (Fig. 4eii). The T4 cells also had small responses to OFF-motion and responded to the ON-like cessation of OFF- motion^25,33^ with responses that are large compared to the T5 responses to the end of ON-motion (Fig. 4eii; UV = 0, Offset response indicated by color change to gray).

Over all UV intensities, there was a change in the mean calcium activity of either the T4 or the T5 ROIs (Fig. 4fi-fii; *t*-test with FDR correction for multiple comparisons, *p* < 0.05, *N*_T4, flies_ = 18, *N*_T5, flies_ = 14). We calculated the isoluminance level of the mean T5 population response (the T5 population isoluminance) as the point when it first reached zero as the UV intensity increased from UV = 0 (Fig. 4fi; green line), and the T4 population isoluminance when the mean response first reached zero as the UV intensity decreased from UV = 15 (Fig. 4fii; purple line). The T4 and T5 population isoluminance levels were 5.2 and 7.8 indicating a substantial difference in their sensitivity to UV. To statistically compare T4 and T5 isoluminance levels, we calculated them for individual ROIs (Fig. 4g; method validated further in Extended Data Fig. 4). The isoluminance levels of T4 ROIs were significantly lower than those of T5 ROIs (Fig. 4h; *p* < 0.001, two-sample *t*-test, *N*_T4, flies_ = 18, *N*_T5, flies_ = 14; *power* = 0.94), and this difference was maintained or strengthened if the ROIs with the strongest responses were selected to evaluate the isoluminance levels (Fig. 4j).

The difference between the T4 and T5 isoluminance levels are consistent with the behavioral responses to ON- and OFF-motion (Figs 1-2), but do not fully explain the full gap between them. While the T4 and T5 population isoluminance levels were 5.2 and 7.8, respectively (Fig. 4h), the ON- and OFF-motion population isoluminance levels were 4.0 and 9.1, respectively (Fig. 2e). Thus, while T4 and T5 are necessary for the behavioral responses (Extended Data Fig. 1g-h, 2c), other cell types may also contribute. In summary, the significantly lower UV-green isoluminance level of T4 cells indicates that they are more sensitive to UV than T5 cells, a difference consistent with the behavioral responses to ON-motion being more sensitive to UV than for OFF-motion.

### Cells presynaptic to T4 have divergent UV-sensitivity consistent with their lamina inputs

To establish how T4 cells are more sensitive to UV than T5 cells, we examined the UV-sensitivity of cells presynaptic to T4. The spectral tunings of cells in the ON-motion pathway are not known, so we systematically measured the responses to expanding UV discs of the major inputs to T4 cells, the Mi1, Tm3, Mi4, Mi9 and C3 cell types^13,26^ (Fig. 5a-b, Extended Data Fig. 5a). We also measured the responses of the lamina monopolar cells (LMCs), the L1, L2, L3, L4 and L5 cell types (Fig. 5a-b), as these cells provide major inputs to the cells presynaptic to T4 and T5^13,26,27^ (Fig. 5c). For a comparison of UV-sensitivity, we also recorded the calcium responses of Dm9, a cell type that is a principal target of R7 photoreceptors^13,36^.

**Figure 5.**
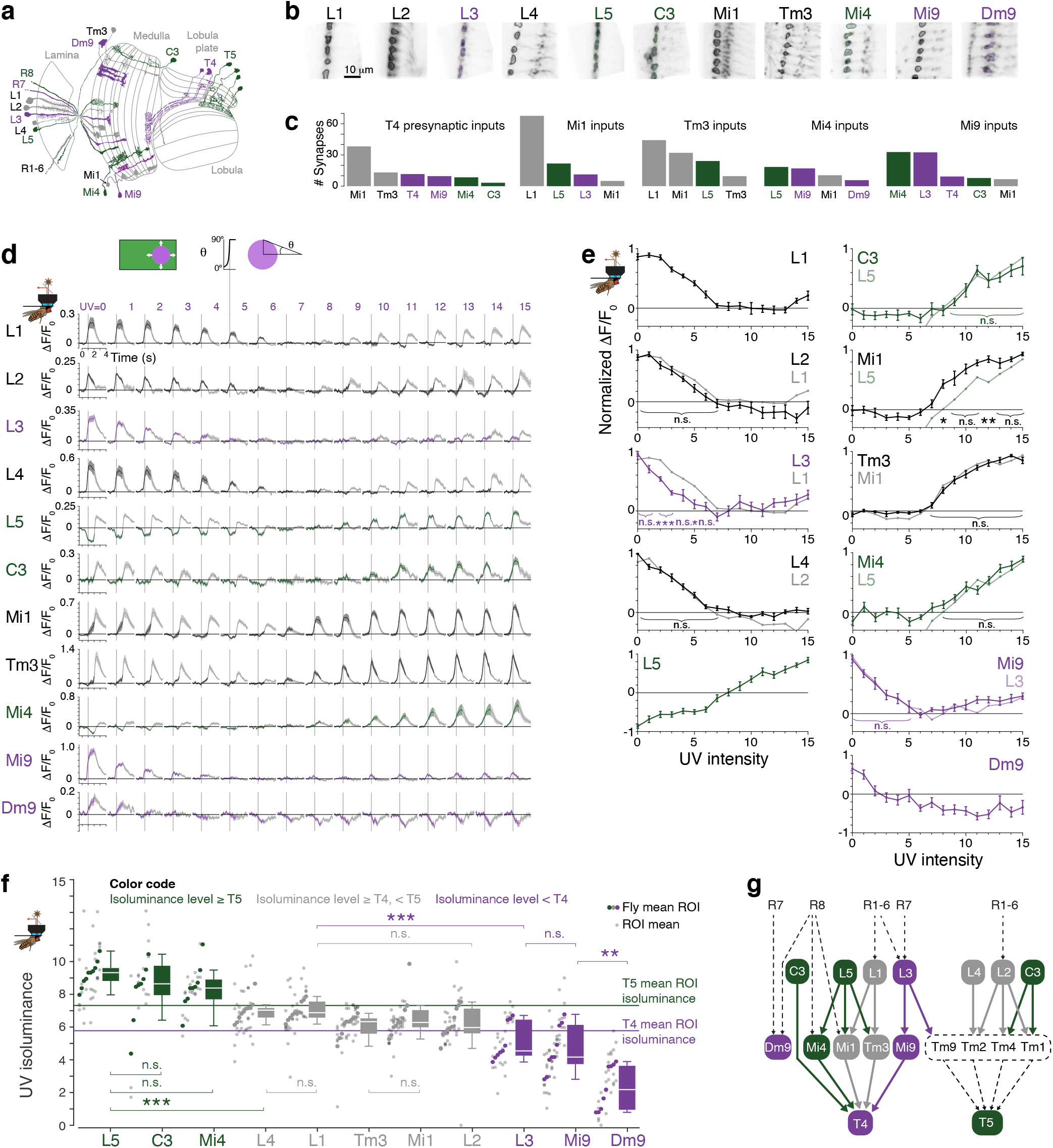
Cells presynaptic to T4 have divergent UV-sensitivity consistent with their lamina inputs. **a**. Diagram of imaged cell types, adapted from^7^. Cell types are color-coded by their isoluminance level (shown in panel f): green indicates an isoluminance level > T5, purple indicates an isoluminance < T4, and gray in-between levels. This color scheme is used throughout the figure. **b**. Examples of ROIs of recorded cell types, with field of view of 34 × 34 μm. Scale bar applies to all images. **c**. Mean number of synaptic inputs to T4 cells from imaged cell types, and of imaged cell types to medulla T4 input cells^26^. The mean number (and percentage) of synaptic inputs from unidentified cells for cell types shown were: 10.9 (9.7%) synapses to T4; 52.2 (20.5%) to Mi1; 29.2 (16.0%) to Tm3; 41.7 (20.5%) to Mi4; 30.4 (17.4%) to Mi9. **d**. Responses of all recorded cell types to expanding UV discs, with the UV intensity indicated above every panel column. Stimulus time course is shown above UV = 5. For every panel, the stimulus starts at *t* = 0, the vertical gray line indicates the end of the disc’s expansion (*t* = 1), whereupon the whole screen remains illuminated by UV for 1 further second (*t* = 2), and then the screen returns to green, indicated by the traces turning to gray. Note that OFF cells have responses to the end of bright discs (e.g., L1, L2, L3, L4, UV = 15, *t* = 2 − 3, responses in gray), and ON cells have responses to the disappearance of dark discs (e.g., L5, C3, Mi1, Tm3, UV = 0, *t* = 2 − 3, responses in gray). Mean ± SEM shown, calculated over flies. *N*_L1,flies_ = 10, *N*_L1,ROI_ = 40; *N*_L2,flies_ = 10, *N*_L2,ROI_ = 39; *N*_L3,flies_ = 9, *N*_L3,ROI_ = 25; *N*_L4,flies_ = 10, *N*_L4,ROI_ = 35; *N*_L5,flies_ = 10, *N*_L5,ROI_ = 30; *N*_C3,flies_ = 9, *N*_C3,ROI_ = 23; *N*_Mi1,flies_ = 10, *N*_Mi1,ROI_ = 36; *N*_Tm3,flies_ = 10, *N*_Tm3,ROI_ = 37; *N*_Mi4,flies_ = 9, *N*_Mi4,ROI_ = 22; *N*_Mi9,flies_ = 9, *N*_Mi9,ROI_ = 41; *N*_Dm9,flies_ = 9, *N*_Dm9,ROI_ = 26; 2-7 ROIs per fly, across cell types. **e**. Mean responses calculated from the mean responses in the 170 ms (two imaging frames) after the disc has expanded (d, vertical gray line in panels at *t* = 1), mean ± SEM over flies shown. Responses of specific cells are overlayed in color, to enable comparisons. For Dm9, responses used for the tuning curve are the first 170 ms (two imaging frames) after the screen has been fully illuminated by UV for 1 s (d, *t* = 2), because the calcium activity dynamics of these cells were much slower than the other cell types we recorded. We used two-sample *t-*tests to compare responses between cell types, with FDR correction for 16 comparisons. Asterisks indicate significance level: *\* p* < 0.05, *\*\* p* < 0.01, n.s. not significant. **f.** Isoluminances of individual ROIs (small, pale gray dots), averaged within each fly (large dots). Boxplot conventions are as in Fig. 2d. Horizontal lines indicate mean ROI isoluminances of T4 and T5, as plotted in Fig. 4h. We tested the hypothesis that the isoluminance levels of pairs of cell types came from the same distribution using two sample *t*-tests, with FDR correction for multiple comparisons. Adjusted *p* values of all comparisons are given in Extended Data Fig. 5c. Asterisks indicate significance level: *\* p* < 0.05, *\*\* p* < 0.01, \*\**\* p* < 0.001, n.s. not significant. Genotypes for all flies used in imaging experiments are in Table 2. **g**. Diagram of connectivity^26,27^ between imaged cells types with cells not imaged in black with dotted lines. Lateral connections within lamina and medulla are not indicated.

All imaged cell types responded robustly to expanding UV discs (Fig. 5d), and we used the same stimulus set as for T4 and T5 cells, discs that expanded to a radius of 30°, to identify responsive ROIs. For the cell types that preferentially responded to OFF-edges, L1-4 and Mi9, we calculated the ROI isoluminance levels using the same methods as for T5 cells. For the cell types that preferentially responded to ON-edges, L5, C3, Mi1, Tm3, and Mi4, we calculated the ROI isoluminance levels using the same methods as for T4 cells.

L1 and L2 are the primary inputs to the ON and OFF-motion pathways, respectively^12,32,37^. Both cell types receive inputs from the R1-6 photoreceptors at shared tetrad synapses and are coupled through gap junctions and chemical synapses in the lamina^23,37^. Because of the close coupling of L1 and L2 cells, we expected them to have similar spectral tunings and in agreement with this prediction their increasing, mean normalized calcium responses were indistinguishable (Fig. 5e; *p* > 0.05, two sample *t*-test with FDR correction for 16 comparisons, *N*_L1, flies_ = *N*_L2,flies_ = 10), although, interestingly, the L2 calcium response decreased at high UV levels (Fig. 5e). The L1 and L2 ROI isoluminance levels were also indistinguishable (Fig. 5f; *p* > 0.5, two-sample *t*-test with FDR correction for 29 comparisons between cell types). The L4 cell type is reciprocally connected to L2 and provides prominent input to the T5 OFF-motion pathway^23,38^. The isoluminance levels of L2 and L4 were also indistinguishable (Fig. 5f; *p* > 0.5, two-sample *t*-test with FDR correction; *N*_L2, flies_ = *N*_L4, flies_ = 10), as were their increasing calcium responses (Fig. 5e; *p* > 0.05, two-sample *t*-test with FDR correction). Thus, closely coupled cells shared similar sensitivities to the UV intensity of expanding discs, providing reassuring evidence for the sensitivity of our measurements.

Two LMCs, L3 and L5, had isoluminance levels that deviated from the shared UV-sensitivity of L1, L2 and L4 (Fig. 5f). L3 receives inputs from the R1-8 photoreceptors^12,13,23^ and provides input to both the T4 and T5 pathways^12,27,39,40^. The isoluminance level of L3 was significantly lower than for the other LMCs, excepting L2 (Fig. 5f; L3-L1 *p* = 0.004; L3-L2 *p* = 0.05; L3-L4 *p* = 0.004; L3-L5 *p* < 0.001; two-sample *t*-test with FDR correction, *N*_L3, flies_ = 9, *N*_L1, flies_ = *N*_L2, flies_ = *N*_L4, flies_ = *N*_L5, flies_ = 10). L5 receives strong input from L1 and L2 in the medulla, from L2 and L4 in the lamina, and provides major input to most of the T4 input neuron types^23,26^. L5 had a response profile very different from the other lamina cells, with a calcium response that decreased at low UV intensities (Fig. 5e). The isoluminance level of L5 was greater than all the other LMCs and also T4 (Fig. 5f; *p* < 0.001, two-sample *t*-tests with FDR correction, *N*_L3, flies_ = 9, *N*_L1, flies_ = *N*_L2, flies_ = *N*_L4, flies_ = *N*_L5, flies_ = 10). We also measured responses in the lamina cell C3 because it provides direct GABAergic, presumed inhibitory, input to T4 cells^26,41^, as well as feedback from the medulla to the lamina where it synapses onto L1, L2, and L3 ^23^. C3 is an ON cell, and the isoluminance levels of L5 and C3 were indistinguishable (Fig. 5f; *p* > 0.05, two-sample *t*-test with FDR correction; *N*_L5, flies_ = 10, *N*_C3, flies_ = 9), as were their increasing calcium responses (Fig. 5e; *p* > 0.05, two-sample *t*-test with FDR correction). These results indicate that lamina cells providing the primary inputs to the motion pathways have UV-green isoluminance levels that differ in their sensitivity to UV, covering a range much broader than the T4-T5 isoluminance difference (Fig. 5f).

We next examined the T4 inputs cells that receive prominent inputs from the L1-5 LMCs. The Mi1 and Tm3 cell types are the principal excitatory inputs to T4 cells^26,41,42^ and they receive major inputs from L1, with a contribution from L5 ^26^ (Fig. 5c). The increasing calcium responses of Mi1 and Tm3 to different intensities of UV discs were not significantly different from each other (Fig. 5e; *p* > 0.05, two-sample *t*-test with FDR correction, *N*_Mi1, flies_ = *N*_Tm3, flies_ = 10), nor were their isoluminance levels (Fig. 5f; *p* > 0.05, two-sample *t*-test with FDR correction). Their isoluminance levels were not significantly different from that of L1, but were from that of L5 (Fig. 5f; Mi1−L1 *p* > 0.05; Tm3−L1 *p* > 0.05; Mi1−L5 *p* < 0.001; Tm3−L5 *p* < 0.001; two-sample *t*-test with FDR correction; *N*_Mi1,flies_ = *N*_Tm3,flies_ = *N*_L1,flies_ = *N*_L5,flies_ = 10), and their increasing calcium responses included significant differences from that of L5 (Fig. 5e; Mi1−L5, UV = 8, 12, *p* < 0.05; Tm3−L5, UV = 8, 12, 14, *p* < 0.05; two-sample t-test with FDR correction). Finally, the isoluminance levels of Mi1 and Tm3 were also not significantly different from T4 (Fig. 5f; Mi1−T4 *p* > 0.05; Tm3−T4 *p* > 0.05; two-sample *t*-test with FDR correction; *N*_Mi1, flies_ = *N*_Tm3, flies_ = 10, *N*_T4, flies_ = 18). These results indicate that Mi1 and Tm3 shared similar tuning to their principal LMC input, L1 and their output target, T4.

The GABAergic Mi4 and glutamatergic Mi9 cell types provide inhibitory inputs to T4^26,41,42^. Based on prior studies, Mi1, Tm3 and Mi4 are ON cells, while Mi9 is unusual for being an OFF cell in the T4 ON-motion pathway^42,43^ — results we have confirmed with our UV discs on a green background (Fig. 5d-e). Mi4 and Mi9 receive their primary LMC inputs from the cells whose isoluminance levels deviated from those of L1 and L2, with L5 the primary LMC input to Mi4, and L3 the primary LMC input to Mi9 (Fig. 5c). The isoluminance level of Mi4 was not significantly different from that of L5 but was significantly greater than that of T4 (Fig. 5f; Mi4-L5 *p* > 0.05, Mi4-T4 *p* < 0.001; two-sample *t*-test with FDR correction; *N*_Mi4, flies_ = 9, *N*_L5, flies_ = 10, *N*_T4, flies_ = 18). Meanwhile, the isoluminance level of Mi9 did not differ from that of L3, significantly less than that of T5, and less but not significantly so than that of T4 (Fig. 5f; Mi9-L3 *p* > 0.05, Mi9-T5 *p* < 0.01; Mi9-T4 *p* > 0.05; two-sample *t*-test with FDR correction; *N*_Mi9, flies_ = 10, *N*_L3, flies_ = 9, *N*_T5,flies_ = 14, *N*_T4,flies_ = 18). These results indicate that Mi4 and Mi9 shared similar tuning to their principal LMC inputs, L5 and L3 respectively, with isoluminance levels significantly outside the values of T4 and T5.

To compare the calcium responses of the cells presynaptic to T4 with those of a cell that receives much of its inputs from R7 cells, we measured the responses to expanding UV discs of Dm9, an UV-OFF cell type and prominent R7 target^13,26,36,44^. Dm9 had a lower UV-green isoluminance level than all the cells in the lamina and T4 motion pathway we recorded, including Mi9 (Fig. 5f; e.g., Dm9–Mi9, *p* = 0.003; two-sample *t*-test, *N*_Mi9, flies_ = 9). Although Dm9 provides a minor input to Mi4 (Fig. 5c), its calcium response was the slowest of the cells we measured (Fig. 5d), indicating it is not likely to drive rapid responses in Mi4.

Together, these results indicate that T4 input cells differ in their sensitivity to UV (Fig. 5f), differences that are consistent with the UV-sensitivity of their primary LMC inputs (Fig. 5g). In particular, L5 drives Mi4, and both cell types have a greater isoluminance level than T4. Complementarily, L3 drives Mi9, and both cell types have lower isoluminance levels than T4 or T5.

### Behavioral responses to UV-Green and Green-UV edges are asymmetric

How does the difference in the UV-sensitivity of the ON- and OFF-motion pathways explain the contribution of color to motion vision? We hypothesized that when the disc is darker than the OFF-motion isoluminance level (UV < 9.2; Fig. 2f), the disc would be dark enough to drive OFF-motion responses, presumably mediated in part by the T5 OFF-motion pathway, as OFF-motion edges did (Fig. 2f). In a complementary way, we hypothesized that when the disc is brighter than the ON-motion isoluminance level (UV > 4.5; Fig. 2e), the disc would be bright enough to drive ON-motion responses, presumably involving the T4 ON-motion pathway, as ON-motion edges did (Fig. 2e). For intensities within this isoluminance band (4.5 < UV < 9.2), UV discs would be both dark enough to generate OFF-motion responses and bright enough to drive the more UV-sensitive ON-motion responses. Based on this hypothesis, we predicted that the fly would respond to an expanding UV disc, whether it is bright, dark, or an intermediate intensity of UV, and so would respond to any intensity of approaching UV discs, consistent with our findings in Fig. 1.

Surprisingly, this hypothesis predicts that the fly’s ability to respond to a moving color edge is affected by its direction of motion (Fig. 6a-b). While the approach of a UV disc on a green background generates motion contrast under our hypothesis, the same is not true for a green disc on a UV background (G-UV), when the green disc is the same intensity as the green background used in our typical experiments, which we refer to here as an intensity-matched green disc. When the background is brighter than the OFF-motion isoluminance (UV > 9.2), we predicted that the intensity-matched green disc is dark enough to drive OFF-motion responses. When the background is darker than the ON-motion isoluminance (UV < 4.5), the disc is bright enough to generate ON-motion responses. But for intermediate intensities (4.5 < UV < 9.2), we predicted that an intensity-matched green disc approaching on a UV background does not generate motion contrast.

**Figure 6.**
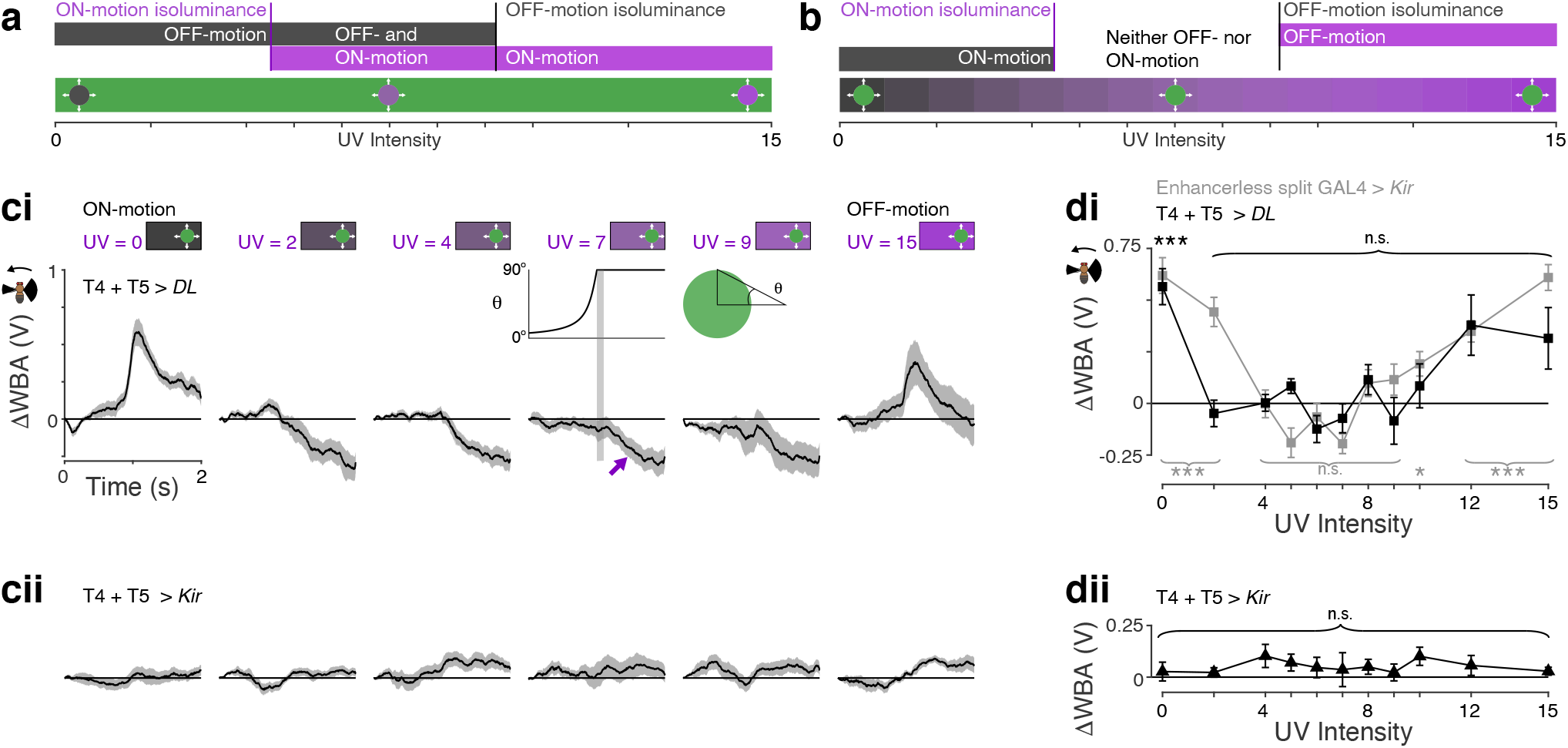
Behavioral responses to UV-Green and Green-UV edges are asymmetric. **a**. Illustration of how a difference in UV-sensitivity of behavioral responses to ON- and OFF-motion enable the detection of isoluminant UV discs expanding on a green background. We hypothesized that when the UV disc is darker than the OFF-motion isoluminance level (UV = 9.2; Fig. 2d), the edge of the disc drives OFF-motion responses (black bar), and when the UV disc is brighter than the ON-motion isoluminance level (UV = 4.5; Fig. 2d), the edge of the disc drives ON-motion responses (purple bar). This hypothesis predicts that when the disc has a UV intensity in the isoluminance band (UV = 4.5 − 9.2), the edge of the disc is simultaneously bright enough (UV > 4.5) to drive ON-motion and dark enough (UV < 9.2) to drive OFF-motion responses (overlap of black and purple bars). **b**. In contrast to panel (a), our hypothesis predicts a lack of responses to a green disc expanding from UV, whose intensity is matched to the green background used in our prior experiments with UV discs. When the UV background is darker than the ON-motion isoluminance level, the edge of the intensity-matched green disc is bright enough to generate ON-motion responses (black bar), and when the UV background is brighter than the OFF-motion isoluminance level, the edge of the disc is dark enough to drive OFF-motion responses (purple bar). When the disc has a UV intensity in the isoluminance band (UV = 4.5 − 9.2), we predicted that the edge of the disc would be neither bright enough (UV > 9) nor dark enough (UV < 9) to drive ON- or OFF-motion responses (gap between black and purple bars). **c**. Turning responses of flies of control genotypes (ci) and T4 and T5 silenced flies (cii) to an intensity-matched green disc expanding on a UV background, with the UV intensity of the background given in purple above panels. The timing of the stimulus is as for the UV discs and shown above the panel for UV = 7. For all genotypes, mean ±SEM shown. **ci**. Wild type *DL* controls for silencing of T4 and T5 cells (black), *N* = 10, and Enhancerless split GAL4 control flies for the expression of *Kir*_2.1_ (gray), *N* = 10. In the panel for UV = 7, the purple arrow indicates the flies turning towards the side the disc appeared from after it has expanded. **cii**. Responses of flies with T4 and T5 cells silenced through expression of *Kir*_2.1_, *N* = 10. **d**. Tuning curves of responses for all UV intensities, for flies shown in (c), measured as the mean response in the 100 ms after the disc has fully expanded, indicated by the gray stripe in the stimulus diagram inset, top row UV = 7. For all rows, *N* =10 flies, and mean ±SEM shown. **di.** To identify responses significantly greater than zero, we used a one tailed *t*-test with FDR correction for 11 comparisons. **dii.** To identify responses significantly different from zero, we used a two-tailed *t*-test with FDR correction for 11 comparisons. Asterisks indicate significance level: *\* p* < 0.05, *\*\* p* < 0.01, \*\**\* p* < 0.001, n.s. not significant. Genotypes for all flies used in behavioral experiments are in Table 1.

In agreement with this stringent prediction, flies did not turn away from an intensity-matched green disc expanding on a UV background over the expected range of UV levels between 4 and 9 (Fig. 6ci, di; T4 + T5 > DL, *p* > 0.05 for 2 ≤ UV ≤ 15; Enhancerless split GAL4 > *kir*, *p* > 0.05 for 4 ≤ UV ≤ 9; one tailed *t*-test that the mean is greater than zero, with FDR correction, *N* = 10). These results revealed an asymmetry in responses to moving color edges, that the response of a fly to an expanding color edge depends on its color polarity, that is whether UV expands into green, or green expands into UV.

To clarify the role of the primary motion pathway in these experiments, we silenced the T4 and T5 cells by expressing *Kir*_2.1_. Under these conditions, flies no longer responded to the direction of the expanding discs (Fig. 6cii, dii; *p* > 0.05, *t*-test that the mean is zero, with FDR correction, *N* = 10). The responses of the control flies indicated that the discs were visible even when the flies didn’t turn away from them. After the discs had fully expanded, the flies reliably turned towards the side the disc came from, for intensities of UV ≥ 2 (Fig. 6ci; *t* = 1 - 2 s, example indicated by a purple arrow for UV = 7). These responses were not just an attraction to a green disc, because they turned away from the discs when the background was dark (Fig. 6ci; UV = 0). Therefore, the turning towards the location of the disc depended on seeing the background UV, and may involve multiple pathways, for example those supporting phototaxis or object vision.

By verifying an unexpected prediction — that green discs do not evoke turning responses over a large range of background UV levels — these experiments support our hypothesis, that a difference in the spectral sensitivity of ON- and OFF-motion underlies the contribution of color to motion vision in flies.

### Enhanced motion detection for approaching objects of selected colors

The mechanisms we have identified in the fly for detecting UV objects could be adjusted to enhance the motion detection of objects of other selected colors too. For example, we expect that the detection of an approaching red object would be enhanced by an augmented red-sensitivity of ON-motion, relative to OFF-motion. To illustrate this hypothesis, we considered the image of an orange in a tree, as seen through a hexagonal lattice of the fly’s compound eye, and estimated the ON- and OFF-motion at every hexagonal pixel as the viewer approached the center of the orange or receded from it (Fig. 7a), by combining vector sums of estimates of the local motion in 4 cardinal directions, mimicking the responses of the ON and OFF pathways (Fig. 7b; see Methods).

**Figure 7.**
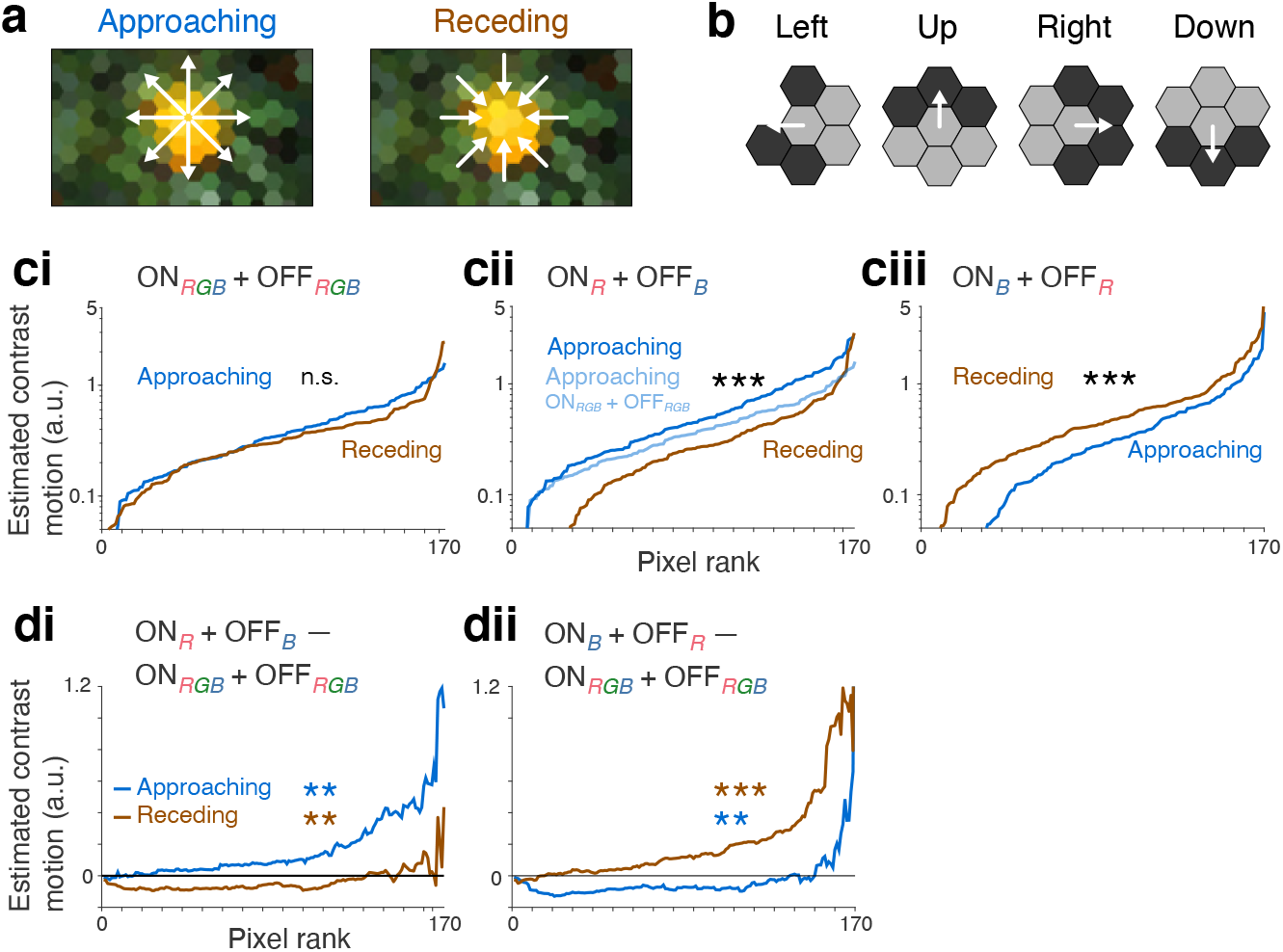
Enhanced motion detection for approaching objects of selected colors. **a.** Photograph of an orange in its tree (*Citrus* sp.), with pixels sampled in a hexagonal lattice of a fly’s eye. Arrows indicate direction of motion for approaching (Left) and receding (Right) from the orange. Photograph: *Kylelovesyou,* Wikimedia Common*s*. **b**. Motion was estimated in four directions using the Weber contrast of pale gray and dark gray pixels; the approaching or receding motion was calculated from their vector sum (see Methods). **c**. Estimated combined ON- and OFF-motion (ON + OFF) at every hexagonal pixel, with pixels rank ordered, for approaching (Blue) and receding (Brown) from the orange, with different combinations of R, G and B intensity values contributing to the estimation of ON- and OFF-motion. **ci**. ON- and OFF-motion (ON*_RGB_* + OFF*_RGB_*) calculated from mean of [R, G, B]. The approaching and receding motion estimates were not significantly different (n.s., *p* > 0.05, Wilcoxon rank sum test, *N_pix_* = 170 hexagonal pixels). **cii**. Red, R, intensity values were used to calculate ON-motion and blue, B, intensity for OFF-motion (ON*_R_* + OFF*_B_*). Approaching and receding motion estimates were significantly different, (***, *p* < 0.001, Wilcoxon rank sum test, *N_pix_* = 170). For comparison, approaching motion calculated using ON*_RGB_* + OFF*_RGB_* from panel (ci) is also shown (Pale blue). **ciii**. B intensity values were used to calculate ON-motion and R intensity for OFF-motion (ON*_B_* + OFF*_R_*). Approaching and receding motion estimates were significantly different, (***, *p* < 0.001, Wilcoxon rank sum test, *N_pix_* = 170). **d**. Comparisons of motion estimates in (c). **di**. Difference between (ON*_R_* + OFF*_B_*) and (ON*_RGB_* + OFF*_RGB_*) estimates. As predicted, changing ON-motion sensitivity to red, compared to OFF-motion, increased the approaching motion (**, *p* < 0.01, Wilcoxon rank sum test, *N_pix_* = 170), and decreased the receding motion (**, *p* < 0.01, Wilcoxon rank sum test, *N_pix_* = 170). **dii**. Also consistent with our predictions, changing ON-motion sensitivity to blue, compared to OFF-motion, decreased the approaching motion (***, *p* < 0.001, Wilcoxon rank sum test, *N_pix_* = 170), and increased the receding motion (***, *p* < 0.01, Wilcoxon rank sum test, *N_pix_* = 170).

When the ON- and OFF-motion are calculated from the same combination of red, green, and blue (R, G, B) intensity values, the magnitude of the approaching and receding motion across the image is nearly identical, as expected (Fig. 7ci; *p* > 0.05, Wilcoxon rank sum test, *N_pix_* = 170 hexagonal pixels). However, in agreement with our hypothesis, when the ON-motion is estimated using the R intensity, and OFF-motion using the B intensity (ON_R_ + OFF_B_), approaching the orange generates greater motion across the image than receding from it (Fig. 7cii; *p* < 0.001, Wilcoxon rank sum test, *N_pix_* = 170), and the motion of approaching is significantly greater than when all R,G,B values are used (Fig. 7di; ON*_RGB_*+OFF*_RGB_* vs ON*_R_*+OFF*_B_*: *p* < 0.01, Wilcoxon rank sum test, *N_pix_* = 170). Conversely, when the color dependency of the ON- and OFF-motion is switched, the estimated receding motion is greater than the approaching motion (Fig. 7ciii; *p* < 0.001, Wilcoxon rank sum test, *N_pix_* = 170), and the receding motion is significantly greater than when all R,G,B values are used (Fig. 7dii; ON*_RGB_*+OFF*_RGB_* vs ON*_B_*+OFF*_R_*: *p* < 0.001, Wilcoxon rank sum test, *N_pix_* = 170).

These results support our hypothesis and demonstrate how motion detection for approaching colored objects in an artificial algorithm can be enhanced by introducing asymmetries in the spectral sensitivity in ON- and OFF-motion detection. The gain in motion detection for the approaching object is tied to a drop in the detection of the receding object, a trade-off that may be acceptable in many situations, for example in automated harvesting systems tailored for specific fruits, or collision avoidance systems.

## Discussion

We have shown that color contributes to motion vision for UV-green edges in *Drosophila* (Fig. 1). Behavioral responses to ON-motion were much more sensitive to UV than responses to OFF-motion (Fig. 2), a difference requiring the R7 photoreceptors (Fig. 3). The T4 and T5 cells that process ON- and OFF-motion, showed a corresponding difference in their sensitivity to UV (Fig. 4), and in the cells linking the R7 photoreceptors and T4 cells, there were consistent spectral differences between lamina monopolar cells and the medulla T4 input cells: L5 and Mi4 were less sensitive to UV, and L3 and Mi9 were more sensitive to UV, compared to the L1 driven Mi1 and Tm3 (Fig. 5g). Finally, we correctly predicted that if the augmented UV-sensitivity of ON-motion processing explained the contribution of color to motion vision, then green discs should not be visible against a UV background that was neither bright nor dark (Fig. 6). We have shown that the contribution of color to motion vision is not just a mechanism for resolving low-contrast UV edges (Fig. 1), but can also be organized to preferentially support the motion detection of objects of specific colors (Fig. 7).

In vertebrates, differences in the spectral sensitivity to ON- and OFF-motion have not been thoroughly investigated, to our knowledge, and if present they could support a contribution of color to motion vision as we have found for *Drosophila*. Larval zebrafish use UV-ON processing to detect paramecia while foraging^45^, and the mechanism we have described has the potential to operate in zebrafish to enhance the detection of approaching paramecia. Mice are also sensitive to UV and green wavelengths, and since they have the greatest chromatic sensitivity in the visual circuits viewing the sky^46^, they are thought to use color vision to detect approaching predators. Again, the mechanism we have described is in theory directly applicable to the mouse visual system: it predicts that if OFF-motion responses were more sensitive to UV than for ON-motion, this would favor the detection of an approaching object seen against a UV-rich sky. In mice, Khani and Gollisch^47^ recently reported ON and OFF retinal ganglion cells that nonlinearly integrate UV and green to allow an OFF cell, for example, to have different isoluminance levels for UV-OFF and green-ON and for green-OFF and UV-ON. In this cell the spectral divergence of ON and OFF processing supports the detection of a light decrement, whether the decrement is in green or UV wavelengths (UV-OFF and green-OFF), and in other cells the nonlinearities were UV-selective (UV-ON and UV-OFF). These nonlinearities are algorithmically very similar to our proposed mechanism and indicate a potential platform for motion-sensitive cells downstream to be sensitive to isoluminant motion, and to preferentially detect objects rich in UV or green, relative to the background. In primates, motion-sensitive cells in area MT contribute to the smooth pursuit tracking of objects and frequently retain some degree of chromatic sensitivity, such that around the isoluminance level the response to motion is decreased, but not to zero^48^. Our results indicate that, in addition to allowing the primates to view isoluminant edges, an individual cell’s chromatic sensitivity may be organized to enhance following the motion of targets of a particular color.

We established that augmented ON-motion UV-sensitivity is not limited to *D. melanogaster* but is also displayed by other drosophilids (Fig. 2g). Among invertebrates, color has been reported to contribute to motion vision in other insects including the honeybee^49^ and the butterfly *Papilio xuthus*^50^, whose behavioral responses to moving colored ON- and OFF-edges indicated that responses to ON-motion were more sensitive to red, compared to responses to OFF-motion that were more sensitive to blue and green. If *Papilio* implements the chromatic mechanism that we have proposed for *Drosophila*, then this would predict that its ON- and OFF-motion pathways support seeing red objects against green backgrounds, for example red flowers set against foliage.

Previous studies have shown that motion vision is colorblind for blue-green gratings in flies^9–11^. We extended that work by not using gratings, which always induce both ON- and OFF-motion, and by developing methods to accurately display wide-field UV-green stimuli. Prior work also indicated that color might contribute to motion vision by broadening the spectral sensitivity of the luminance channel through unidentified cellular mechanisms^14^, and subsequent EM reconstructions indicated that the R7 and R8 photoreceptors form synapses in the medulla with cells specifically presynaptic to T4^12,26,27^. Our results indicate that UV-sensitivity is maintained along the R7-L3-Mi9-T4 pathway (Fig. 5g), and predict that R7 cells innervate L3. Indeed, we recently demonstrated in an EM reconstruction study that R7 cells form substantial numbers of previously unreported synapses with L3 and other cells in the optic chiasm between the lamina and medulla^13^. To understand how sensitivity to UV propagates from R7 through the lamina and medulla circuitry to T4 cells (Figs 4 and 5) is complex due to sign changes, asymmetric spectral tuning, and recurrent connections along the pathway. To focus on just one example, the Mi4 and Mi9 cell types, which are inhibitory to T4 cells^42^, heavily synapse onto each other^26^ and reciprocal inhibition between these cell types may amplify their chromatic differences.

In summary, we have shown how UV contributes to the processing of ON-motion in *Drosophila* in a way that enhances responses to expanding UV discs. We have identified key cellular components of how color contributes to motion vision in flies, the R7 and T4 cells, and how cells linking them show consistent differences in their spectral tuning. We have shown how a spectral divergence in ON- and OFF-motion processing can be used to favor objects of a specific color, an insight that is directly applicable to many vertebrate and invertebrate sensory systems.

## Acknowledgements

We thank Na Ji for suggesting Teflon® for the projection screen; Emily Behrmann and David Stern lab for assistance with *Drosophila* species; Eyal Gruntman and members of the Reiser lab, Ben Hardcastle, and Hiroshi Shiozaki for comments on the manuscript; and Mikko Juusola, Bevil Conway, and the many Janelia colleagues for helpful discussions. This work is funded by HHMI.

## Methods

### Contact for reagent and resource sharing

Further information and requests for resources and reagents should be directed to and will be fulfilled by Michael Reiser.

### Fly Stocks

All flies were reared on a standard cornmeal and agar diet. Flies were kept at 21°C and 60% humidity on a 16 hour ON: 8 hours OFF light cycle prior to behavior and imaging experiments. All *D. melanogast*er used for behavior and imaging contained at least one copy of the wild type *white* allele to ensure completely wild type eye pigmentation, and *w*^+^{*DL*} indicates *white* alleles from the Dickinson Lab strain, a strain generated from 200 wild caught flies in the lab of Michael Dickinson, Caltech, Pasadena, CA, USA.

We used split GAL4 lines characterized in our previous work: the lamina cell types lines are as used in ^51^, the T4/T5 and the medulla T4 input cell lines are as used ^42^, and Dm9 line is as in ^41^. The exception is the split GAL4 line used to image L1, shown in Extended Data Fig. 5a, which is improved from the L1 split-GAL4 lines used in ^51^. The genotypes used for behavior experiments are listed below in Table 1, and those used for imaging are listed in Table 2. For all behavioral results, all the primary data were from enhancerless split GAL4 crossed with wild type *DL* flies unless otherwise stated. The enhancerless split GAL-4 flies have transgenes in the same genomic location as the other split-GAL4 drivers, and so match the general genotype, but these transgenes lack the enhancer-containing cis-regulatory sequences that determine the specific patterns of the other driver lines.

**Table 1.** Genotypes of flies used in behavior experiments.

| Genotype | Rescued photoreceptors | Neurons manipulated | Fig. panel |
| --- | --- | --- | --- |
| *w+ norpA<sup>36</sup>/ w+ norpA<sup>36</sup>; UAS-norpA/+; Rh1-Gal4(attP2)/+* | R1-R6 | R7 R8 Ocelli | 1di 1ei 3a-b |
| *w+ norpA<sup>36</sup>/ w+ (DL); UAS-norpA/+; Rh1-Gal4(attP2)/+* | All | None (R7 R8 Ocelli control) | 1dl 1ei 3a-b |
| *w+(DL)/w<sup>1118</sup>; BPp65ADZp(attP40)/ +(DL); BPZpGdbd(attP2)/ +(DL)* | N/A | N/A (Enhancerless split GAL4 control, ES > DL) | 1diii 1eiii 2b-g<br>3c Ex.1f<br>Ex.2a Ex.3c<br>Ex.4a-b |
| *Drosophila mauritiana* | N/A | N/A | 2g |
| *Drosophila sechellia* | N/A | N/A | 2g |
| *Drosophila santomea* | N/A | N/A | 2g |
| *Drosophila yakuba* | N/A | N/A | 2g |
| *Drosophila teissieri* | N/A | N/A | 2g |
| *w+ norpA<sup>36</sup>/ w+ norpA<sup>36</sup>; UAS-norpA/+; Rh1-Gal4(attP2)/ Rh3-Gal4(attP2)* | R1-R6 pR7 | yR7 R8 Ocelli | 3a, Ex.3a |
| *w+ norpA<sup>36</sup>/ w+ (DL); UAS-norpA/+; Rh1-Gal4(attP2)/ Rh3-Gal4(attP2)* | All | None (yR7 R8 Ocelli control) | 3a, Ex.3a |
| *w+ norpA<sup>36</sup>/ w+ norpA<sup>36</sup>; UAS-norpA/+; Rh1-Gal4(attP2)/ Rh4-Gal4(attP2)* | R1-R6 yR7 | pR7 R8 Ocelli | 3a, Ex.3a |
| *w+ norpA<sup>36</sup>/ w+ (DL); UAS-norpA/+; Rh1-Gal4(attP2)/ Rh4-Gal4(attP2)* | All | None (pR7 R8 Ocelli control) | 3a, Ex.3a |
| *w+ norpA<sup>36</sup>/ w+ norpA<sup>36</sup>; UAS-norpA/+; Rh1-Gal4(attP2)/ Rh5-Gal4(attP2)* | R1-R6 pR8 | R7 yR8 Ocelli | 3a, Ex.3a |
| *w+ norpA<sup>36</sup>/ w+ (DL); UAS-norpA/+; Rh1-Gal4(attP2)/ Rh5-Gal4(attP2)* | All | None (R7 yR8 Ocelli control) | 3a, Ex.3a |
| *w+ norpA<sup>36</sup>/ w+ norpA<sup>36</sup>; UAS-norpA/+; Rh1-Gal4(attP2)/ Rh6-Gal4(attP2)* | R1-R6, yR8 | R7 pR8 Ocelli | 3a, Ex.3a |
| *w+ norpA<sup>36</sup>/ w+ (DL); UAS-norpA/+; Rh1-Gal4(attP2)/ Rh6-Gal4(attP2)* | All | None (R7 pR8 Ocelli control) | 3a, Ex.3a |
| *w+ norpA<sup>36</sup>/ w+ norpA<sup>36</sup>; UAS-norpA/+; Rh1-Gal4(attP2)/Rh2-Gal4(attP2)* | R1-R6 Ocelli | R7 R8 | 3a, Ex.3a |
| *w+ norpA<sup>36</sup>/ w+ (DL); UAS-norpA/+; Rh1-Gal4(attP2)/Rh2-Gal4(attP2)* | All | None (R7 R8 control) | 3a, Ex.3a |
| *w+ norpA<sup>36</sup>/ w+ norpA<sup>36</sup>; UAS-norpA/ Rh3-Gal4(BDSC# 7457); Rh1-Gal4(attP2)/ Rh5Gal4(attP2)* | R1-R6 pR7 pR8 | yR7 yR8 Ocelli | 3a, Ex.3a |
| *w+ norpA<sup>36</sup>/ w+ (DL); UAS-norpA/ Rh3-Gal4(BDSC# 7457); Rh1-Gal4(attP2)/Rh5-Gal4(attP2)* | All | None<br>(yR7 yR8 Ocelli control) | 3a, Ex.3a |
| *w+ norpA<sup>36</sup>/ w+ norpA<sup>36</sup>; UAS-norpA/ Rh3-Gal4(BDSC# 7457); Rh1-Gal4(attP2)/Rh4-Gal4(attP2)* | R1-R6 R7 | R8 Ocelli | 3a, Ex.3a |
| *w+ norpA<sup>36</sup>/ w+ (DL); UAS-norpA/Rh3-Gal4(BDSC# 7457); Rh1-Gal4(attP2)/Rh4-Gal4(attP2)* | All | None<br>(R8 Ocelli control) | 3a, Ex.3a |
| *w+ norpA<sup>36</sup>/ w+ norpA<sup>36</sup>; UAS-norpA/Rh6-Gal4(BDSC# 7459); Rh1-Gal4(attP2)/Rh4-Gal4(attP2)* | R1-R6 yR7 yR8 | pR7 pR8 Ocelli | 3a, Ex.3a |
| *w+ norpA<sup>36</sup>/ w+ (DL); UAS-norpA/Rh6-Gal4(BDSC# 7459); Rh1-Gal4(attP2)/Rh4-Gal4(attP2)* | All | None (pR7 pR8 Ocelli control) | 3a, Ex.3a |
| *w+ norpA<sup>36</sup>/ w+ norpA<sup>36</sup>; UAS-norpA/Rh6-Gal4(BDSC# 7459); Rh1-Gal4(attP2)/Rh5-Gal4(attP2)* | R1-R6 R8 | R7 Ocelli | 3a, Ex.3a |
| *w+ norpA<sup>36</sup>/ w+ (DL); UAS-norpA/Rh6-Gal4(BDSC# 7459); Rh1-Gal4(attP2)/Rh5-Gal4(attP2)* | All | None<br>(R7 Ocelli control) | 3a, Ex.3a |
| *w+(DL)/w<sup>1118</sup>;*<br>*59E08-p65ADZp(attP40)/tubP-GAL80<sup>ts</sup>;*<br>*42F06-ZpGdbd(attP2)/ UAS-Kir2.1* | N/A | T4 T5<br>(T4 T5 > Kir) | 6cii 6dii<br>Ex.1gi Ex.1hi<br>Ex.2b-c |
| *w+(DL)/w<sup>1118</sup>;*<br>*59E08-p65ADZp(attP40)/+(DL);*<br>*42F06-ZpGdbd(attP2)/ +(DL)* | N/A | None<br>(T4 T5 > DL) | 6ci 6di<br>Ex.1gii Ex.1hii<br>Ex.2b-c |
| *w+(DL)/w<sup>1118</sup>;*<br>*BPp65ADZp(attP40)/ tubP-GAL80<sup>ts</sup>;*<br>*BPZpGdbd(attP2)/ UAS-Kir2.1* | N/A | None (Enhancerless split GAL4 Kir2 control, ES > Kir) | 6di Ex.1hii |
| *w+(DL)/w<sup>1118</sup>; +/+(DL);*<br>*Rh1-Gal4(attP2)/UAS-shibire<sup>ts</sup>* | N/A | R1-R6<br>(Rh1 > shi) | Ex.3b |
| *w+(DL)/w<sup>1118</sup>; +/+(DL);*<br>*Rh1-Gal4(attP2)/ +(DL)* | N/A | None (R1-6 control, Rh1 > DL) | Ex.3b |
| *w+(DL)/w<sup>1118</sup>; +/+(DL);*<br>*Rh3-Gal4(attP2)/ UAS-shibire<sup>ts</sup>* | N/A | pR7<br>(Rh3 > shi) | 3c Ex.3c |
| *w+(DL)/w<sup>1118</sup>; +/+(DL);*<br>*Rh3-Gal4(attP2)/ +(DL)* | N/A | None (pR7 control, Rh3 > DL) | 3c Ex.3c |
| *w+(DL)/w<sup>1118</sup>; +/+ (DL); Rh4-Gal4(attP2)/ UAS-shibire<sup>ts</sup>* | N/A | yR7<br>(*Rh4* > *shi*) | 3c Ex.3c |
| *w+(DL)/w<sup>1118</sup>; +/+ (DL); Rh4-Gal4(attP2)/ +(DL)* | N/A | None (yR7 control,<br>*Rh4* > *DL*) | 3c Ex.3c |
| *w+(DL)/w<sup>1118</sup>; +/+ (DL); Rh5-Gal4(attP2)/ UAS-shibire<sup>ts</sup>* | N/A | pR8<br>(*Rh5* > *shi*) | 3c Ex.3c |
| *w+(DL)/w<sup>1118</sup>; +/+ (DL); Rh5-Gal4(attP2)/ +(DL)* | N/A | None (pR8 control,<br>*Rh5* > *DL*) | 3c Ex.3c |
| *w+(DL)/w<sup>1118</sup>; +/+ (DL); Rh6-Gal4(attP2)/ UAS-shibire<sup>ts</sup>* | N/A | yR8<br>(*Rh6* > *shi*) | 3c Ex.3c |
| *w+(DL)/w<sup>1118</sup>; +/+ (DL); Rh6-Gal4(attP2)/ +(DL)* | N/A | None (yR8 control,<br>*Rh6* > *DL*) | 3c Ex.3c |
| *w+(DL)/w<sup>1118</sup>; +/+ (DL); Gal4(attP2)/ UAS-shibire<sup>ts</sup>* | N/A | None (Enhancerless<br>GAL4 > *shi*, EG > *shi*) | Ex.3b |
| *w+(DL)/w<sup>1118</sup>; +/+ (DL); Gal4(attP2)/ +(DL)* | N/A | None (Enhancerless<br>GAL4 > *DL*, EG > *DL*) | Ex.3b |

**Table 2.** Genotypes of flies used in imaging experiments.

| **Genotype** | **Cells Targeted** | **Indicator** | **Fig. Panel** |
| --- | --- | --- | --- |
| *w+(DL)/w<sup>1118</sup>; 59E08-p65ADZp(attP40)/+; 42F06-ZpGdbd(attP2)/ 20XUAS-IVS-NES-jRGECO1a-p10 (VK00005)* | T4/T5 | jRGECO1a | 4c-j 5f<br>Ex.4a-b |
| *w+(DL)/w<sup>1118</sup>; 40F12-p65ADZp(attP40)/+; VT027316-ZpGdbd(attP2)/ 20XUAS-IVS-NES-jRGECO1a-p10 (VK00005)* | L1 | jRGECO1a | 5b 5d-f<br>Ex.5b-c |
| *w+(DL)/w<sup>1118</sup>; 53G02-p65ADZp(attP40)/+; 29G11-ZpGdbd(attP2)/ 20XUAS-IVS-NES-jRGECO1a-p10 (VK00005)* | L2 | jRGECO1a | 5b 5d-f<br>Ex.5b-c |
| *w+(DL)/w<sup>1118</sup>; 59A05-p65ADZp(attP40)/+; 75H07-ZpGdbd(attP2)/ 20XUAS-IVS-NES-jRGECO1a-p10 (VK00005)* | L3 | jRGECO1a | 5b 5d-f<br>Ex.5b-c |
| *w+(DL)/w<sup>1118</sup>; 31C06-p65ADZp(attP40)/+; 34G07-ZpGdbd(attP2)/ 20XUAS-IVS-NES-jRGECO1a-p10 (VK00005)* | L4 | jRGECO1a | 5b 5d-f<br>Ex.5b-c |
| *w+(DL)/w<sup>1118</sup>; 21A05-p65ADZp(attP40)/+; 31H09-ZpGdbd(attP2)/ 20XUAS-IVS-NES-jRGECO1a-p10 (VK00005)* | L5 | jRGECO1a | 5b 5d-f<br>Ex.5b-c |
| *w+(DL)/w<sup>1118</sup>; 26H02-p65ADZp(attP40)/+; 29C11-ZpGdbd(attP2)/ 20XUAS-IVS-NES-jRGECO1a-p10 (VK00005)* | C3 | jRGECO1a | 5b 5d-f<br>Ex.5bc |
| *w+(DL)/w<sup>1118</sup>; 55C05-p65ADZp(attP40)/+; 71D01-ZpGdbd(attP2)/ 20XUAS-IVS-NES-jRGECO1a-p10 (VK00005)* | Mi1 | jRGECO1a | 5b 5d-f<br>Ex.5b-c |
| *w+(DL)/w<sup>1118</sup>; 38C11-p65ADZp(attP40)/+; 59C10-ZpGdbd(attP2)/ 20XUAS-IVS-NES-jRGECO1a-p10 (VK00005)* | Tm3 | jRGECO1a | 5b 5d-f<br>Ex.5b-c |
| *w+(DL)/w<sup>1118</sup>; 48A07-p65ADZp(attP40)/+; 79H02_ZpGdbd(attP2)/ 20XUAS-IVS-NES-jRGECO1a-p10 (VK00005)* | Mi4 | jRGECO1a | 5b 5d-f<br>Ex.5b-c |
| *w+(DL)/w<sup>1118</sup>; 48A07-p65ADZp(attP40)/+; VT046779-ZpGdbd(attP2)/20XUAS-IVS-NES-jRGECO1a-p10 (VK00005)* | Mi9 | jRGECO1a | 5b 5d-f<br>Ex.5b-c |
| *w+(DL)/w<sup>1118</sup>; 19G04-p65ADZp(attP40)/+; 53A05-ZpGdbd(attP2)/20XUAS-IVS-NES-jRGECO1a-p10 (VK00005)* | Dm9 | jRGECO1a | 5b 5d-f<br>Ex.5b-c |
| *w<sup>1118</sup>; 40F12-p65ADZp(attP40)/+; VT027316-ZpGdbd(attP2)/pJFRC51-3XUAS-IVS-Syt::smHA (su(Hw)attP1), pJFRC225-5XUAS-IVS-myr::smFLAG (VK00005)* | L1 | HA and<br>FLAG | Ex.5a |

Rh3- and Rh6-GAL4 lines with insertions on the second chromosome were obtained from the Bloomington stock center (BDSC #7457 and #7459, respectively). To generate additional Rh1-, Rh2-, Rh3-, Rh4-, Rh5- and Rh6-GAL4 driver lines with the transgenes inserted in the attP2 landing site on the third chromosome, we PCR-amplified previously characterized promoter regions^52–57^ from genomic DNA, TOPO-cloned the PCR products into pENTR-D-TOPO and transferred to pBPGUw (Addgene #17575) using standard Gateway cloning. Primer sequences were as described^13^ (Rh3, Rh5, Rh6) or as listed below (Rh1, Rh2, Rh4). Transgenic flies were generated by phiC31-mediated integration (Genetic Services, Inc.).

> Rh1F CACC GGC ATT GAC ACA TTA AAT CGC TG
>
> Rh1R TCA CTG GGG CGG ACT AGT CGC
>
> Rh2F CACC TTC TGG CTG CCC TTT AGT GTC A
>
> Rh2R GCT CAG CTA CCC GCA ACC CCT T
>
> Rh4F CACCTT GAA CCG ATG TGG CAG CAC CA
>
> Rh4R TTC GAA TGG CTG GTA CTG GTG

### Immunohistochemistry

To visualize the expression pattern of the L1 split-GAL4 driver line (Extended Data Fig. 5a), we used pJFRC51-3XUAS-IVS-Syt::smHA in su(Hw)attP1 and pJFRC225-5XUAS-IVS-myr::smFLAG in VK00005 ^58^. The images were generated by the Janelia FlyLight project team. Sample preparation was as previously described^59^ and a full protocol is available online, https://www.janelia.org/project-team/flylight/protocols under “IHC - Anti-GFP”, “IHC - Polarity Sequential” and “DPX mounting”. Images were acquired on Zeiss LSM 710 or 800 confocal microscopes with 20x 0.8 NA or 63x 1.4 NA objectives. For display, we generated resampled views from three-dimensional image stacks using the Neuron annotator mode of V3D ^60^ and exported these images as TIFF format screenshots.

### Visual display and calibration

We displayed stimuli using customized DLP Lightcrafter projectors (v2, Texas Instruments Inc., Dallas, TX, USA). We replaced the blue LED with a 385 nm 1650 mW UV LED (item # M385D2, Thorlabs Inc., Newton, NJ, USA), inserted a bandpass filter in front of the green LED (item # FF01-554/23-21.8-D, Semrock Inc., Rochester, NY, USA), and disconnected the red LED. The plastic diffusers and lenses are thin enough to pass UV with little attenuation. Stimuli were displayed at 120 Hz, with a frame update rate of 60 Hz, a pixel resolution of 608 x 684 pixels, and the maximum 4-bit color depth, so color intensities ranged 0−15, limiting the UV intensities levels to the range 0−15.

UV-green patterns were rear-projected onto a projection screen of Teflon film (item # 8569K, McMaster-Carr Supply Co., Chicago, IL, USA). UV and green wavelengths are scattered differently by the screen, and as the effect is large (Extended Data Fig. 1a), it is imperative that this is corrected in a UV display system. To correct for this, we created a gimbal from two manual rotation stages (MSRP01, Thor Labs, Newton, NJ, USA) that allowed us to measure the irradiance at the precise location of the fly’s head of the visual display in 10° steps, comprising 10 x 25 measurements (model USB4000-UV-VIS, with QP600-2-UV-VIS light guide, Ocean Optics Inc., now Ocean Insight, Largo, FL, USA). We created a luminance mask for the green channel that adjusted the green light intensity at every location to match UV light intensity (Extended Data Fig. 1a-e), adjusted by a constant linear scaling factor, which we set to be 2.3 after iterative sets of behavioral experiments so that the isoluminant UV intensity had a mid-range value roughly in the middle of the intensity range of 0 and 15. As a result, the green illumination pattern is fixed in all experiments where there is green light. The UV intensity varies slightly across the screen (Extended Data Fig. 1a), and as we could not create a luminance mask for the UV-channel and maintain the ability to change the UV intensity, the UV intensity is expressed by the intensity value (0-15) and not the irradiance (but we note that the ratio of UV to Green at each location is tightly controlled after the calibration, Extended Data Fig. 1e). The effectiveness of this approach was validated by the motion isoluminance shown by colorblind *norpA*^36^ mutants with *norpA* function restored in R1-6 using *Rh1*-GAL4 (Fig. 1d-e, 3b).

Two projector systems were used to collect the behavioral data, created and calibrated identically. All the data presented in the main figures were collected on the same projector system. Data from the second system is used for Extended Data Fig. 2a only.

For the imaging experiments, one projector was created and calibrated as for the behavior experiments. To minimize the spectral overlap between the display’s illumination and the sensitivity of the detection pathway of the 2-photon microscope, we made 2 modifications: a filter with a narrower pass band was used in front of the green LED (Item # FF01-549/12-25-D, Semrock Inc.) and additional short pass filter was placed in front of the projector lens (item # SP01-561RY-25, Semrock Inc.). The display spanned −20° to 100° azimuth and −50° to +50° elevation, from the perspective of the average mounted fly.

### Behavioral measurements

We cold anaesthetized 2–5-day old female flies and glued them to a 0.1 mm tungsten rod (catalog # 71600; A-M Systems) on the dorsal prothorax for positioning, using UV-curing glue (KOA300-1, KEMXERT). They recovered for at least 1 hour prior to tests.

The tethered fly was illuminated from above by an infrared LED and the amplitudes of the shadows of its wing beats were monitored by an optical wing-beat analyzer, which consists of optical sensors connected to custom hardware. The difference in the amplitudes of the shadows of the wingbeats (ΔWBA) measured the turning response, and we sampled it at 500 Hz using a data acquisition card (NI PCI-6221, National Instruments) and data acquisition toolbox in MATLAB.

For flies expressing *shibire*^ts1^ and several control genotypes (Fig. 3c, Extended Data Fig. 3b-c), we exposed the flies to the specified temperatures indicated in each figure panel, by placing them in a temperature-controlled incubator, with a humidity of 60% and white-light illumination, for 40 minutes prior to an experiment. Individual experiments then lasted 25 minutes and were conducted at 21°C.

For painting the ocelli, we used black oil paint (Winsor and Newton, Artist’s Oil Colour 386), and sealed the paint with a thin coat of UV-curing glue.

### Visual stimuli for behavioral experiments

We created and controlled visual stimuli using the Psychophysics Toolbox^61^ in MATLAB (Mathworks, Natick, MA, USA), and a Nvidia GeForce GTX 770 2GB GDDR5 PCI Express 3.0 graphics card (Nvidia Corp., Santa Clara, CA, USA). We organized stimuli into trials of 8 s duration. The first 6 s were closed-loop stripe fixation, with the fly’s turning response (ΔWBA) controlling the position of a 10° azimuth wide black bar, moving on a green background. The stimulus was presented in the last 2 s of the trial. Two types of stimuli were used, expanding discs and competing ON- and OFF-motion, described below. In total, four protocols were used: 1) expanding UV discs with a green background; 2) expanding Green discs with a UV background; 3) expanding UV discs with a green background of different speeds; 4) competing ON-motion edges over the range UV = 0 – 15; 5) competing OFF-motion over the range UV = 0 – 15; 6) competing ON- and OFF-motion over the range UV = 3-9.

### Expanding UV discs with a green background

For flies viewing UV discs expanding from a green background (Fig. 1d-e, Extended Data Fig. 1g-h), the disc appeared at 6 s and expanded as though moving towards the fly with a constant velocity until it had expanded to fill the visual display after one more second, at 7 s. The size of a disc expanding with apparent constant motion (displayed with an accelerating angular size) can be parameterized by the ratio of its radius, *r*, to its velocity, *v*, and for all experiments *r*/*v* = 120 ms, except where stated. The expanded disc remained on the screen for the last 1 s, so for a UV = 15 disc, the display remained UV = 15 during this period. Expanding discs were presented on both sides, either at +60° azimuth, elevation 0°, or at −60° azimuth, elevation 0°, and the responses averaged with the response inverted for the left-hand responses.

In every set of trials, expanding discs were shown with UV intensities of {0, 2, 4, 5, 6, 7, 8, 9 10, 12, 15}. We also measured responses to a blank green screen, a blank UV = 15 screen, and to black and green square wave gratings with a spatial wavelength of 30°, and a temporal frequency of 5 and 10 Hz, for clockwise and anticlockwise yaw rotations, to generate optomotor responses (data not shown). All stimuli were presented 5 times (10 times including from the left or right, including blank screens). The stimulus presentation order was randomized for each set of trials. The protocol took 20 minutes to complete.

### Expanding green discs with a UV background

For flies viewing green discs expanding from a UV background (Fig. 6c-d, the screen switched from green to UV at 6s, and the green disc appeared. The disc then expanded to full size after one more second, at 7 s, and the screen remained green for another second, until 8 s. The disc expanded with *r*/*v* = 120 ms, and at ±60° azimuth, 0° elevation as for the UV expanding discs. The UV intensity of the background varied, but the calibrated green intensity of the disc was fixed.

In every set of trials, expanding green discs were shown with UV intensities of the background of {0, 2, 4, 5, 6, 7, 8, 9 10, 12, 15}. In all other respects, the stimuli were organized as for the expanding UV discs.

### Expanding UV discs with a green background of different speeds

For the measurements of response to discs expanding with different apparent speeds (Extended Data Fig. 1f), the expansion speed was varied for UV = 6 discs appearing out of a green background for *r*/*v* of {10, 15, 30, 60, 120} ms. We also measured responses to black and green square wave gratings, spatial wavelength 30°, temporal frequency of 5 and 10 Hz, for clockwise and anticlockwise yaw rotations, to generate optomotor responses (data not shown). All stimuli were presented 5 times (10 times including from the left or right), and the stimulus presentation order was randomized for each set of trials.

### *Competing ON-motion* over the range UV = 0 – 15

During the competing ON-motion stimuli that covered the range of UV intensities from 0 to 15 (Fig. 2b-e), the screen switched from green to black (unilluminated) at 6 s. The visual display was split into 8 windows of 30° azimuth, perspective-corrected so that they were of equal angular extent. For clockwise stimuli, a green edge appeared at 6 s from the left-hand side of every window and moved rightwards at 120°s^-1^ to fill the window by 250 ms (Fig. 2a). Simultaneously a UV edge appeared at 6 s on the right-hand side of every window and moved leftwards to fill the window by 250 ms. This sequence repeated every 250 ms, a temporal frequency of 4 Hz, and lasted for 2 s (8 cycles total). Counterclockwise stimuli were presented in the same way but reflected along a vertical axis so that green edges moved leftwards, and UV edges moved rightwards. The green intensity was fixed, and the UV intensities were {0, 2, 4, 5, 6, 7, 8, 9 10, 12, 15}.

As for the expanding UV discs, we also measured responses to black and green square wave gratings, spatial wavelength 30°, temporal frequency of 5 and 10 Hz, for clockwise and anticlockwise yaw rotations, to generate optomotor responses. All stimuli were presented 5 times (10 times including from the left or right), and the stimulus presentation order was randomized for each set of trials.

### *Competing OFF-motion* over the range UV = 0 – 15

During the competing OFF-motion stimuli that covered the range of UV intensities from 0 to 15 (Fig. 2b, d, f), the same frames were displayed as used for the competing ON-motion stimuli but shown in the reverse temporal order (Fig. 2a). After 6 s, the screen switched to all green and UV. For clockwise stimuli, a green edge receded from the left-hand side of every window, retreating rightwards to void the window of green by 250 ms. Simultaneously, a UV edge receded from the right-hand side of every window and retreated leftwards to void the window of UV, leaving the screen blank by 250 ms. As for ON-motion, counterclockwise stimuli were presented in the same way, but reflected along a vertical axis so that green edges moved leftwards, and UV edges moved rightwards. In all other respects, the stimuli were organized as for competing ON-motion.

### Competing ON- and OFF-motion over the range UV = 3 – 9

To measure responses of flies to both ON and OFF-motion (Figs 2g, 3, Extended Data Figs 2, 3), we presented UV intensities over the limited range {3, 4, 5, 6, 7, 8, 9}, so that responses to both competing ON and competing OFF-motion could be measured in the same flies within a protocol that took 21 minutes. We also measured responses to black and green square wave gratings, spatial wavelength 30°, temporal frequency of 5 and 10 Hz, for clockwise and anticlockwise yaw rotations, to generate optomotor responses. All stimuli were presented 5 times (10 times including from the left or right), and the stimulus presentation order was randomized for each set of trials.

### Data analysis for behavioral experiments

All data were analyzed in MATLAB. The responses to clockwise and counterclockwise stimuli were averaged, with the counterclockwise responses inverted. Likewise, responses to expanding discs centered on opposing azimuth locations of ±60° were inverted and combined.

For the responses to expanding discs, the response in the 100 ms after the disc had expanded was used to calculate how the response varied with UV intensity. For the competing edge stimuli, the mean response over the duration of the stimulus (2 s) was used to calculate how the response varied with UV intensity.

The isoluminance level was calculated as the first point when the fly’s mean response over all trials was greater than zero, as the UV intensity increased from UV = 0, using linear interpolation between stimulus intensities, as illustrated in Fig. 2e-f.

For statistical tests of responses to expanding discs, we used student’s *t*-test to assess if the response were significantly different from zero (Fig. 1e, Extended Data Fig. 1g-h, 2c), and tested its normality using the Anderson-Darling test. We controlled for multiple comparisons by applying a false discovery rate (FDR) correction to a significance level of 0.05. For all these experiments there were at least 10 flies, except for the measurements of experiments with different speeds (Extended Data Fig. 1f) where there were 7 flies.

For competing motion over the range UV = 0 – 15, we compared responses to ON- and OFF-motion using two-sample student’s *t*-test to compare responses (Fig. 2d) and tested its normality using the Anderson-Darling test. For these experiments there were 10 flies for both conditions.

For experiments with competing ON- and OFF-motion over the range UV = 3 – 9, the isoluminance levels were capped at UV = 3 and UV = 9 because of the restricted range of the stimuli. For these responses, we used non-parametric methods, the Wilcoxon signed rank test for paired samples, and Wilcoxon rank sum test for two independent samples. To control for multiple comparisons between genotypes we applied an FDR correction to a significance level of 0.05. For all experiments there were at least 10 flies.

### Calcium Imaging

We cold anaesthetized 2–5-day old female flies and glued a metal pin to the thorax for positioning, using a UV-curing glue that dries to a rigid consistency (Loctite 3972). We cut off the front legs and the proboscis and covered the wounds with a UV-curing glue with less UV absorption and reemission than others, Bondic®. We tilted the head forward so that the dorsal part of the back of the head was horizontal and glued the head and prothorax to the aperture of the imaging shim and immersed in saline solution: 103mM NaCl, 3mM KCl, 1.5mM CaCl_2_, 4mM MgCl_2_, 26mM NaHCO_3_, 1mM NaH_2_PO_4_, 8mM trehalose, 10mM glucose, 5mM TES. The cuticle over the right-hand side of the back of the head was perforated and removed using a fine insect pin bent into a fine hook, in the style of the dental scaler tool, in combination with forceps. The air sacks, fat cells and trachea were removed so that the medulla, lobula and lobula plate were accessible for imaging. Experiments were performed at 21° C.

Cells were imaged using a two-photon microscope (Prairie, Ultima; Bruker) with infrared excitation (1060 nm, Coherent Chameleon Ultra II) delivering less than 21 mW power at the sample, and a x40 objective (Nikon 40X CFI APO NIR). We imaged T4 and T5 cells at 6.83 Hz, 256 x 256 pixels, x16 zoom, with a field of view of 17 x 17 μm. We imaged all other cells at 6.147 Hz, 256 x 256 pixels, and x8 zoom, with a field of view of 34 x 34 μm. Responses were recorded for 7.6 s of every trial, and the data saved in the last 0.4 s of every trial. Before the experimental protocol was displayed, expanding discs expanding up to θ = 30° were shown repeatedly to center the imaging field of view to the same region of visual space across flies and experiments.

### Visual stimuli for calcium imaging

We presented expanding UV discs with a green background using the same dynamics and spatial location as used for the behavioral experiments. The discs were centered on azimuth −60° elevation 0°, and the disc expanded at t = 0 from θ = 6.8° with *r*/*v* = 120 ms to full expansion (θ = 90°) after 1 s, followed by the screen remaining illuminated by the fully expanded disc for 1 s, before resetting to green. For 3s before and after the stimulus, the screen was green, and the trials lasted 8s in total. Flies also viewed directional green and black ON- and OFF-motion edges, with a spatial wavelength of 30° and a temporal frequency of 1 Hz, moving up, down, left, or right (Fig. 4c-d). For the ON-motion stimuli, the screen switched to blank and then green edges appeared, as for the green component of the competing motion stimuli in behavioral experiments. We also recorded responses to a uniform green screen, a uniform UV = 15 screen, and an unilluminated, blank screen (data not shown). All stimuli were shown for 5 trials, with the order of stimuli randomized for every set of trials, taking 18 minutes in total.

For the experiments with expanding UV discs with a UV background (Extended Data Fig. 4), the stimuli were as for the expanding UV discs with a UV background, except that the background was set to UV = 5 for T4 cells, and the background was set to UV = 8 for T5 cells. Before and after the stimuli, the screen was green, and in all other regards, the trials were organized exactly as for the expanding UV discs with a green background.

### Data analysis for calcium imaging

Imaging data was recorded for the first 7.6s of every trial, that lasted 8s, and the frames were saved during the last 0.4s. The visual display was triggered by the onset of image recording, and the stimulus frames were temporally synchronized to the image acquisition.

To spatially align frames, we calculated a binary template of the mean calcium fluorescence for one stimulus (UV = 0 for OFF cells, or UV = 15 for ON cells) and calculated the spatial cross-correlation between the template and binarized frames, for all frames in the experiment. Recordings with large movement (>25 pixels) were discarded.

To calculate regions of interest (ROIs), we took the mean response to one stimulus (UV = 0 or UV = 15), over the duration of recording (7.6 s), spatially smoothed the signal with a Gaussian filter with a full width at half maximum of 11 pixels and identified all the peaks in the calcium fluorescence. We then used a flood filling algorithm to identify the ROIs around these peaks, creating ROIs with separations that corresponded to medulla column widths of ∼5 μm, and lobula column widths of ∼4 μm (T4 and T5 example ROIs in Fig. 4c-d; examples of ROIs for other cell types in Fig. 5b). To calculate the relative change in fluorescence, ΔF/F_0_, we calculated the initial fluorescence, F_0_, for every stimulus trial as the fluorescence in the 1.5 s before the stimulus is displayed.

The semi-automated ROI detection process produced between 2 and 5 ROIs per fly for T4 recordings, and between 3 and 6 ROIs per fly for T5 recordings. These ROIs are based on mean calcium levels and included ROIs with very small responses to expanding discs. We excluded visually unresponsive ROIs by applying a threshold for the peak responses. For ON-sensitive cells, the threshold was applied to the mean response to UV expanding discs over the range UV = 12-15. For OFF-sensitive cells, the threshold was applied to the mean response to expanding UV discs over the range UV = 0-3. For Dm9, which is an OFF cell with a very restricted range of responses exhibiting calcium increases in our setup, the threshold was applied to the mean response to expanding UV discs over the range UV = 0-1.

For the T4 cells, this procedure identified 26% (16/62) of ROI as unresponsive, and for T5 cells, it identified 41% (24/58) of ROIs as unresponsive. The effects of the thresholding procedure on the isoluminance level are shown in Fig. 4j for T4 and T5, and in Extended Data Fig. 5b for other cell types. As the threshold increases, the isoluminance estimate remains stable and the number of flies decrease. We chose values that eliminated non-responding ROIs, as indicated by the gray vertical lines in Fig. 4j and Extended Data Fig. 5b.

For all cells we used the two imaging frames immediately after the disc has fully expanded to characterize how the response changed with UV intensity, corresponding to an interval of 150 ms for T4 and T5, and an interval of 170 ms for the other cell types. For Dm9, we used the first 170 ms after the screen has switched back to green, as the cells continue to respond to the 1 s display of UV after the disc has expanded, and this gave a more reliable estimate of their isoluminance levels. For the responses of T4 and T5 ROIs to ON- and OFF-motion edge stimuli, we calculated as the mean response during the stimulus (lasting 2 s). For the ON-motion stimuli, the mean response to the screen switching to black before the edges appear was subtracted from the stimulus responses.

For all ON-sensitive cells (T4, L5, C3, Mi1, Tm3, and Mi4), the ROI isoluminance level was the first point when the response is less than zero using linear interpolation between stimulus intensities, as the UV intensity is decreased from UV = 15 (illustrated for T4 in Fig. 5g). For all OFF-sensitive cells (T5, L1, L2, L3, L4, Mi9, and Dm9) the isoluminance level was the first point when the response was less than zero using linear interpolation between stimulus intensities, as the UV intensity is increased from UV = 0 (illustrated for T5 in Fig. 4g). In the rare cases when there was no zero crossing (6/46 T4 ROIs, 1/34 T5 ROIs for the data in Fig. 4), the minimum was chosen. For every fly, we calculated a mean isoluminance of that fly’s ROIs, and we calculated the overall mean ROI isoluminance as the average over flies, so that individual flies contributed equally to the population statistics (Figs 4h, 5f). As an additional method to average out noise in responses, we also calculated the population isoluminance, the UV intensity when the mean response across flies, first reached zero (using the same ON/OFF consideration as above; used in Fig. 4f). In this calculation, every fly again contributed equally to the mean response.

To statistically test the differences between the isoluminances of cell types, we calculated the mean isoluminance across ROIs for individual flies and used the student’s *t*-test, after checking the data was normally distributed, using the Anderson-Darling test. To control for multiple comparisons between cell types (Fig. 5f), we used FDR correction. In Fig. 5f, there were 29 comparisons, whose results are all displayed in Extended Data Fig. 5c. For all imaging data, the number of samples is given as the number of flies, *N*_flies_.

We verified our methods for measuring the isoluminance levels using UV discs expanding from a UV background (Extended Data Fig. 4a). For T4 recordings, the background was UV = 5, setting the isoluminance level to be unambiguously UV = 5, and indeed we measured a mean ROI isoluminance level of 5.2 ±0.3 (mean ±SEM, *N*_flies_ = 9; Extended Data Fig. 4b). For T5 recordings, the background was UV = 8, setting the isoluminance level to be UV = 8, and we measured a mean ROI isoluminance level of 7.7 ±0.3 for T5 (mean ±SEM, *N*_flies_ = 8; Extended Data Fig. 4b), which was also in good agreement with the predetermined value.

### Data analysis of photographic images

For Fig. 7, we converted the original 900 x 1500 pixel image into a hexagonal photograph of 170 hexagons, each 120 pixels wide and 104 pixels high, by replacing the RGB intensity values within a hexagon with the mean values of the 9360 pixels that hexagon. At every hexagonal facet, we estimated the motion in the left, up, right, and down directions. To do this, we calculated the average intensity of the home and neighboring facets, *I_h_*, (pale gray hexagons in Fig. 7b), and the average intensity of background neighboring facets along the direction of motion, *I_b_*, (dark gray hexagons in Fig. 7b), and calculated the Weber contrast as C = (*I_h_* - *I_b_*)/ *I_b_*. The approaching ON-motion was then calculated as the vector sum of positive Weber contrast along the direction of the hexagon relative to the point of expansion, and the approaching OFF-motion was calculated as the vector sum of negative Weber contrast along the direction of the hexagon relative to the point of expansion. Likewise, the receding ON-motion was the vector sum of positive Weber contrast along the direction of the hexagon relative to the point of contraction and receding OFF-motion was the vector sum of negative Weber contrast along the direction of the hexagon relative to the point of expansion. These calculations are instantaneous estimates of motion that neglect, for example, differences in depth caused by a change in viewing position but serve to illustrate the mechanism explored here. The ON- and OFF-motion estimates were then combined with equal weight (ON + OFF in Fig. 7), to form the approaching and receding motion values across the hexagons of the image. This process was repeated using the mean RGB values for both ON- and OFF-motion (ON_RGB_ + OFF_RGB_), using R values for ON- and B values for OFF-motion (ON_R_ + OFF_B_), and using R values for ON- and B values for OFF-motion (ON_B_ + OFF_R_).

## Extended Data Figure Legends

**Extended Data Figure 1.**
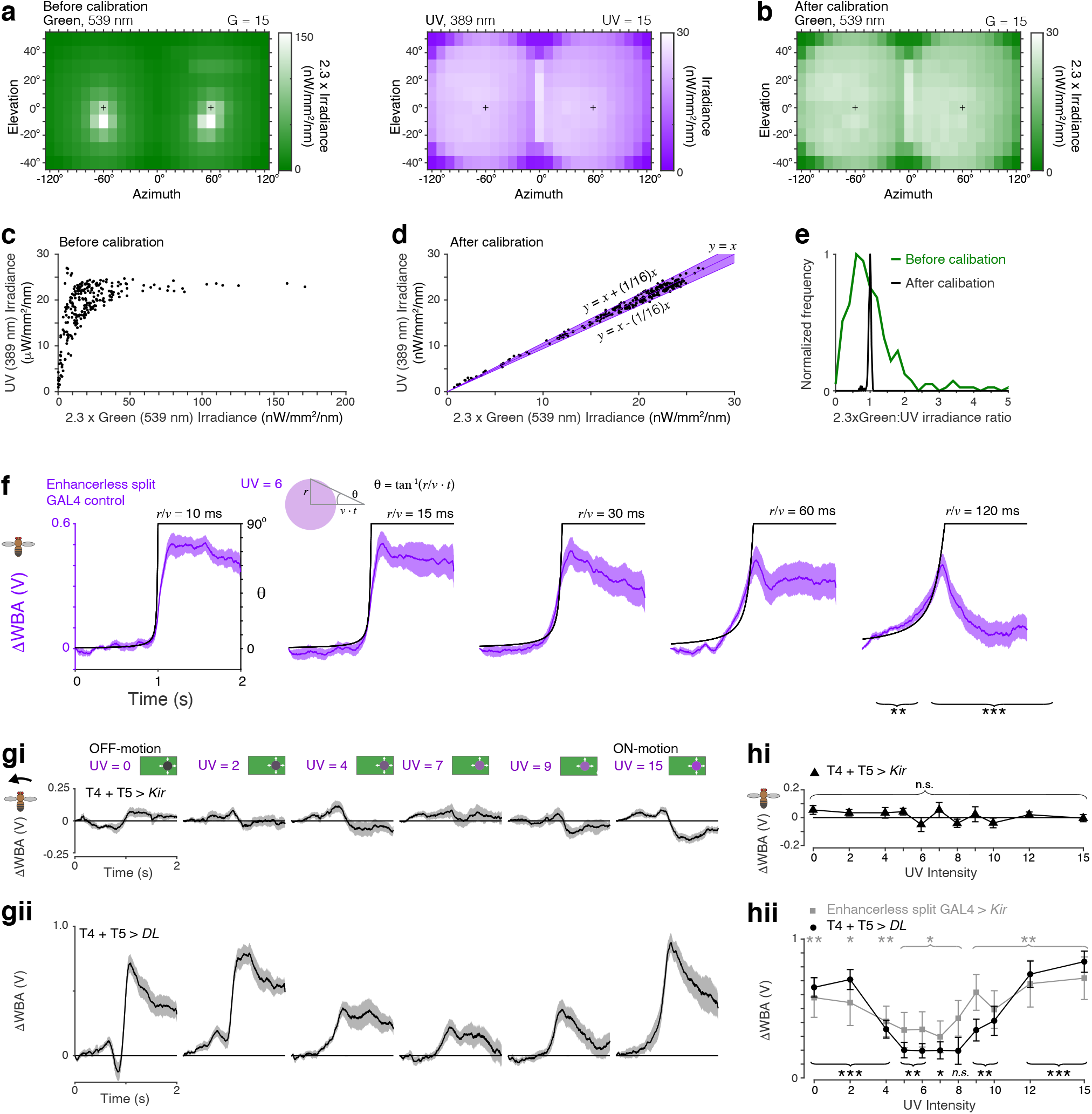
Calibration of display system, speed tuning of the contribution of color to motion vision, and requirement of T4 and T5 cells. **a**. Spatial distributions across the display screens of the irradiance of the peak green (539 nm, left) and UV (389 nm, right) wavelengths prior to calibration. For details of the calibration, see Methods. The green irradiance was calibrated to a scaled factor of 2.3, relative to the UV, as this resulted in behavior with isoluminance for *norpA*^36^ Rh1-rescue flies in the mid-range of UV intensities. Black crosses at azimuth ±60° elevation 0° indicate the foci of the expanding discs. **b**. Spatial distributions of the scaled irradiance of the peak green wavelength after calibration. **c, d.** Distribution of all scaled green irradiance measurements plotted against the UV irradiance, prior to calibration in (c), and after calibration in (d). The irradiances were matched to the precision of the color bit depth, 1/16, indicated by the purple band. **e**. Histograms of the ratio of the green irradiance scaled by 2.3 and the UV irradiance, for every location measured before and after calibration, corresponding to the points plotted in panels (c) and (d). **f**. Responses of enhancerless split GAL4 control flies to approaching UV discs of different speeds, defined by *r*/*v*, with a UV intensity of 6. Black lines indicate the angle, θ, subtended by the radius of the disc. Mean ±SEM responses shown, *N* = 12. **g**. Responses of flies with both T4 and T5 silenced by expression of *Kir*_2.1_ (top, **gi**) compared to genetic control flies (bottom, **gii**). Mean ±SEM responses shown, *N* = 10. Turning responses are abolished by the expression of *Kir*_2.1_ in T4 and T5 for all intensities of UV, quantified in (h). **h**. Tuning curves of turning responses in (g), along with an additional control genotype, enhancerless split GAL4 > *Kir*_2.1_, measured as the mean response in the first 100 ms after the disc has fully expanded. **hi**. Mean turning responses are not significantly different from zero with the expression of *Kir*_2.1_ in T4 and T5 (*p* > 0.05; *t*-test with FDR correction for multiple comparisons, *N* = 10). **hii**. Mean turning responses of control significantly different from zero with the expression of *Kir*_2.1_ in T4 and T5 (*p* > 0.05; *t*-test with FDR correction for multiple comparisons, *N* = 10). Genotypes of all flies used in behavioral experiments are listed in Table 1.

**Extended Data Figure 2.**
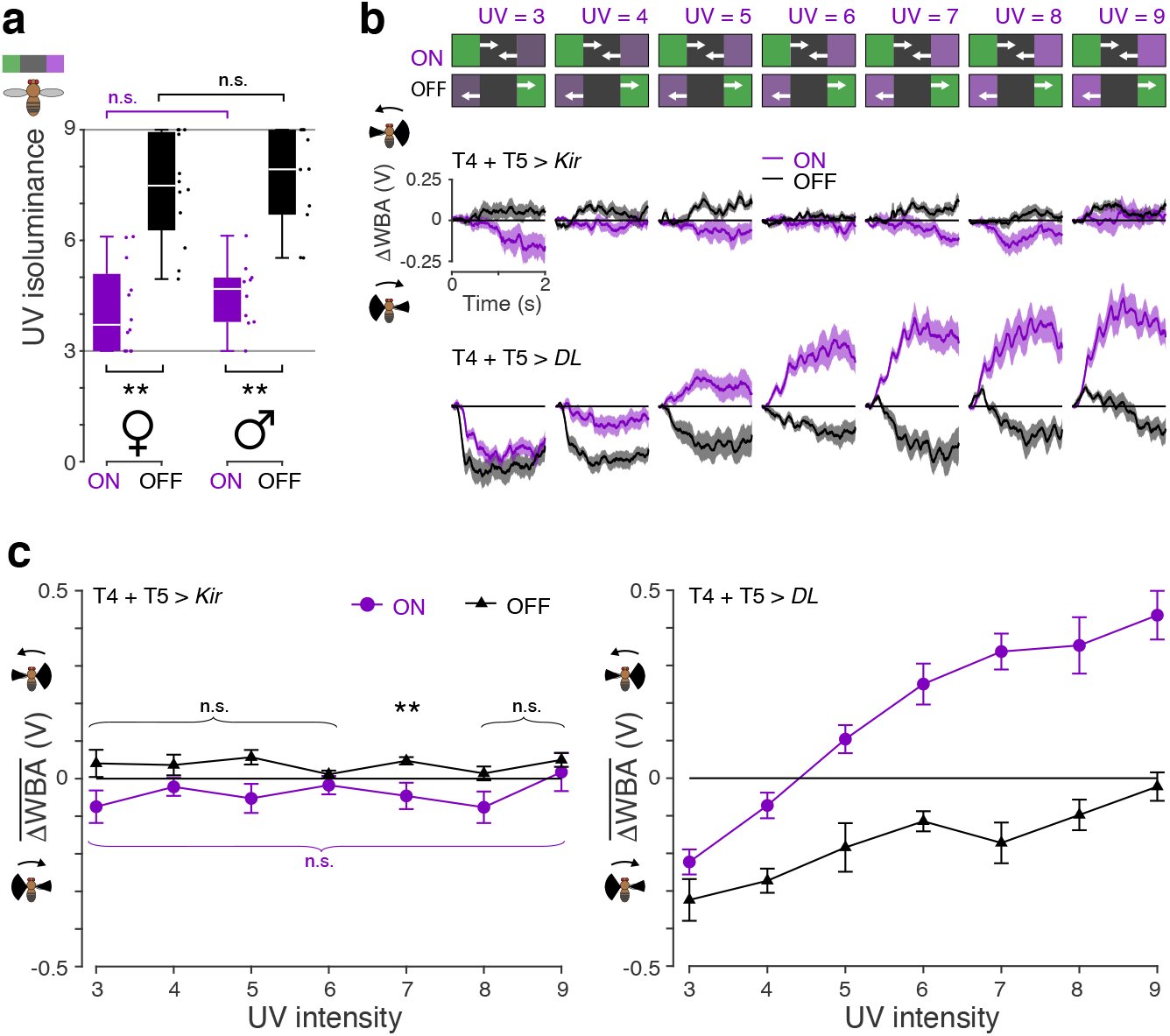
Male and female behavioral responses to competing ON- and OFF-motion, and the requirement of T4 and T5 cells. **a**. Competing ON- and OFF-motion isoluminance levels of female (left) and male flies (right) measured in a replica setup of the one used to collect the behavioral data for all the main figures. Neither the competing ON- nor the competing OFF-motion isoluminance levels were significantly different between the sexes (*p* ≥ 0.3, Wilcoxon signed rank test, *N* = 10), but were within each sex (*p* ≤ 0.003, Wilcoxon signed rank test, *N* = 10). Boxplot conventions are as in Fig. 2d. **b.** The direction-selective T4 and T5 neurons are required for responses to competing ON- (purple) and OFF-motion (black). Responses of flies with T4 and T5 silenced by expression of *Kir*_2.1_ (top) and genetic control flies (bottom). To measure responses to competing ON- and OFF-motion in the same flies, the UV intensities were restricted to the range UV = 3 − 9. Mean ±SEM responses shown, *N* = 10. **c**. Mean responses to competing ON- and OFF-motion of all flies with T4 and T5 silenced by expression of *Kir*_2.1_ (left) and genetic control flies (right). Mean ±SEM responses shown, NT_4T5>*Kir*_ =10, *N*_T4T5>*DL*_ =10. Mean turning responses are abolished by the expression of *Kir*_2.1_ in T4 and T5 (student’s *t*-test was used to identify responses significantly different from zero, with FDR correction for 7 comparisons; asterisks indicate significance: *\*\* p* < 0.01). The exception was for UV = 7 for competing OFF-motion, where the response magnitude was small at 0.048 but significant (*p* = 0.006). Genotypes of all flies used behavioral experiments are listed in Table 1.

**Extended Data Figure 3.**
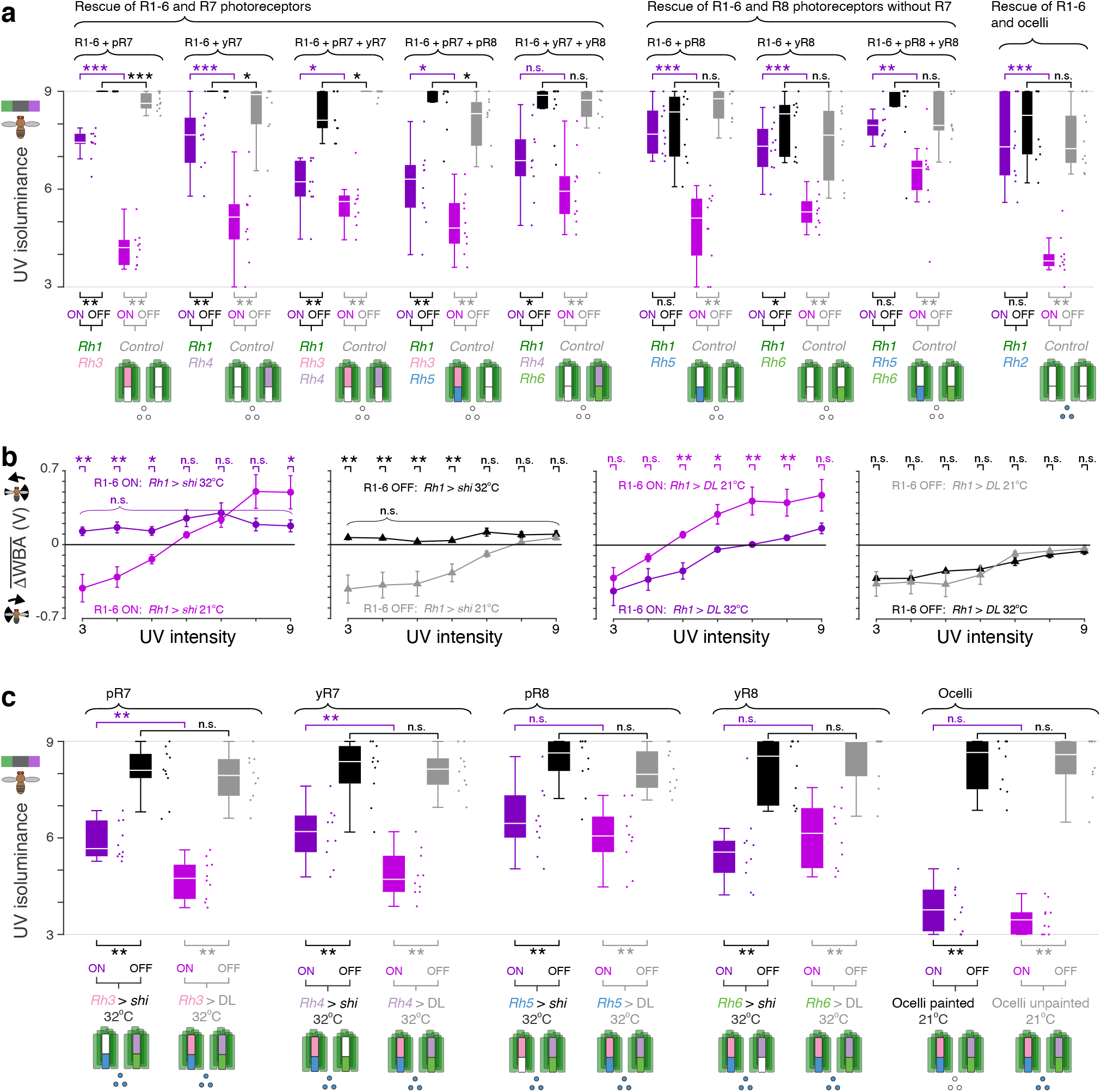
Isoluminance levels for ON- and OFF-motion for genetic rescue and silencing experiments. **a**. Isoluminance levels for competing ON- (purple) and OFF-motion (black) of homozygous *norpA*^36^ flies with the function of different combinations of photoreceptors rescued using rhodopsin-GAL4 driven expression of UAS-*norpA*, along with isoluminance levels for competing ON- (light purple) and OFF-motion (gray) of genetic controls. The UV intensity was restricted to the range 3-9 to enable both isoluminance levels to be measured in the same flies — the gray horizontal lines at UV = 3 and 9 indicate these bounds. Below each pair of boxplots, for every rescue genotype and control, a diagram visually indicates the photoreceptors rescued (colored) and not rescued (white). We used paired Wilcoxon signed rank test to compare isoluminance levels within genotypes, and two sample Wilcoxon rank sum tests to compare isoluminance levels between rescue and control genotypes, with *N* = 10 for all genotypes. Asterisks indicate significance level: *\* p* < 0.05, *\*\* p* < 0.01, \*\**\* p* < 0.001, n.s. not significant. Boxplot conventions are as in Fig. 2d. **b**. We silenced R1-6 using *Rh1*-GAL4 expression of UAS-*shibire*^ts1^ and heating the flies to 32°C. Plots show mean ±SEM turning response over the 2 s of the competing motion stimuli for genotypes at 32°C and 21°C. We used two sample Wilcoxon rank sum tests with a FDR correction for multiple comparisons to compare responses at specific UV intensities between temperature conditions, with *N* = 10 for all genotypes and temperature conditions. Leftmost panel: responses of *Rh1* > *shi* flies to competing ON-motion silenced at 32°C (purple) and at room temperature, 21°C (light purple). At the restrictive temperature, 32°C, there was no significant difference in the responses from the mean (*p* > 0.05, *t*-test, FDR correction, *N* = 10). Middle left panel: responses of *Rh1* > *shi* flies to competing OFF-motion silenced at 32°C (black) and at room temperature, 21°C (gray). At the restrictive temperature, 32°C, there was no significant difference in the responses from the mean (*p* > 0.05, *t*-test, FDR correction, *N* = 10). Middle right panel: responses of control *Rh1* > wild type *DL* flies to competing ON-motion at 32°C (purple) and at room temperature, 21°C (light purple). For the darkest (UV = 3) and brightest (UV = 9) tested intensities of UV, there was no significant difference in the responses at the two temperatures, however, the increased temperature increased the isoluminance level of the mean response. Rightmost panel: responses of control *Rh1* > wild type *DL* flies to competing OFF-motion at 32°C (black) and at room temperature, 21°C (gray). There was no significant difference in the responses at the two temperatures for all tested UV intensities. **c**. Isoluminance levels for competing ON- (purple) and OFF-motion (black) of flies with UAS-*shibire*^ts1^ expressed in different classes of photoreceptors through rhodopsin GAL4 driver lines, and in an enhancerless GAL4 control, along with isoluminance levels for competing ON- (light purple) and OFF-motion (gray) of genetic controls (based on same data as in Fig. 3c). The UV intensity was restricted to the range 3-9 to enable both isoluminance levels to be measured in the same flies — the gray horizontal lines at UV = 3 and 9 indicate these bounds. Below each pair of boxplots, for every rescue genotype and control, diagram visually indicates the photoreceptors silenced (white) and not silenced (colored). We used paired Wilcoxon signed rank test to compare isoluminance levels within genotypes, and two sample Wilcoxon rank sum tests to compare isoluminance levels between rescue and control genotypes, with *N* = 10 for all genotypes. Plotting conventions for asterisks as in panel (a). Boxplot conventions are as in Fig. 2d. Genotypes for all flies used in behavioral experiments are in Table 1.

**Extended Data Figure 4.**
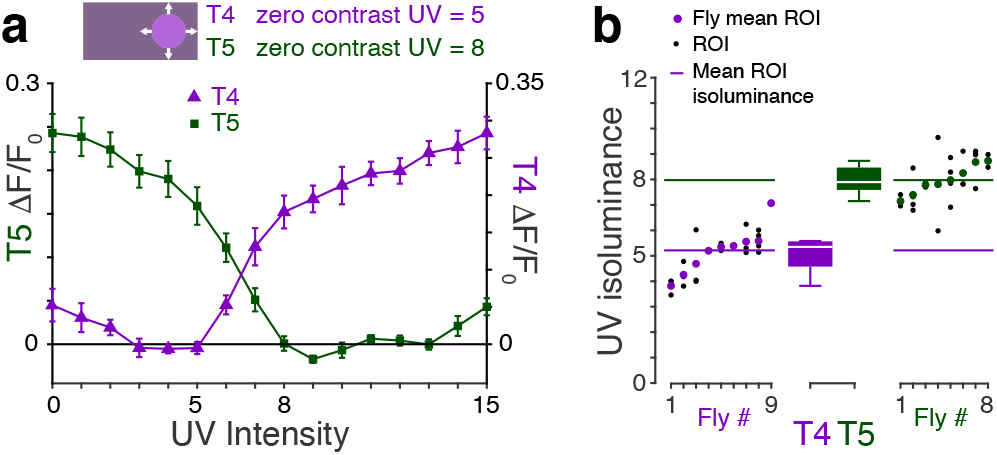
Validation of isoluminance determination for T4 and T5 using contrast tuning. **a**. Mean calcium activity responses of T4 and T5 ROIs to UV discs expanding out of a UV background. For the T4 recordings, the background UV intensity was UV = 5, and so when the discs shared this intensity, they were unambiguously isoluminant. For the T5 recordings, the background UV intensity was UV =8, and so when the discs shared this intensity, they were unambiguously isoluminant. We chose these background intensities to test our methods at comparable illuminance levels to the values determined from the experiments with UV discs expanding out of a green background (Fig. 4f). Mean ±SEM shown, *N*_T4, flies_ = 9, *N*_T5, flies_ = 8, different flies to those in Fig. 4. **b**. Isoluminance levels of ROIs (black dots) and flies (colored circles) for T4 (purple) and T5 (green) cells. Boxplot conventions are as in Fig. 2d. Colored horizontal lines indicate the mean ROI isoluminances: 5.2 ±0.3 for T4 (mean ±SEM, *N*_T4, flies_ = 9), and 7.7 ±0.3 for T5 (mean ±SEM, *N*_T5, flies_ = 8).

**Extended Data Figure 5.**
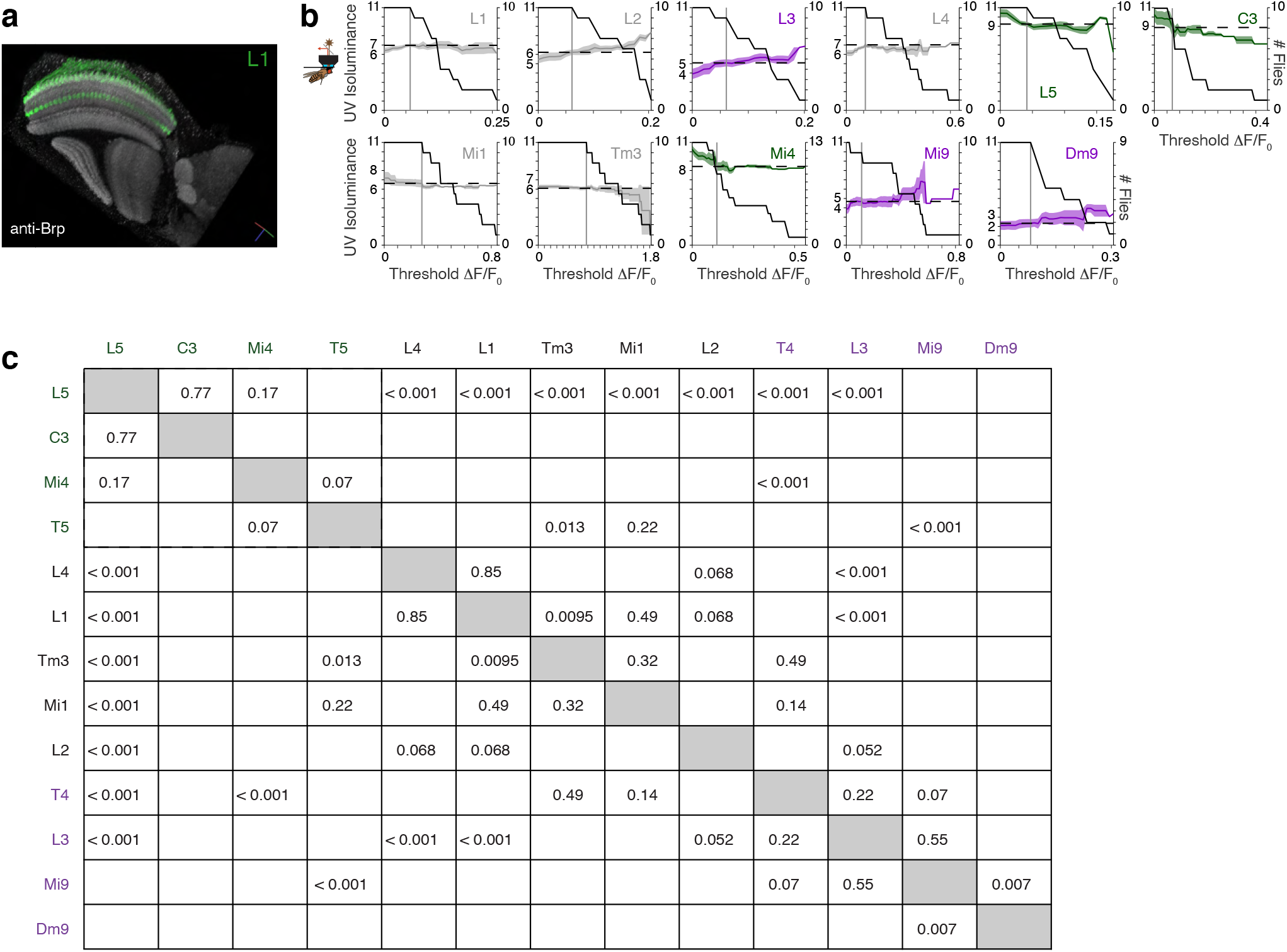
Imaging of T4 input cells, lamina cells, and Dm9. **a**. Horizontal section through the optic lobe visualizing the expression pattern (green) for the L1 split GAL4 line showing distinguishing expression in medulla layers 1 and 5 (see Fig. 5a). Gray is nc82 antibody staining for Bruchpilot to indicate neuropils. **b**. Mean ROI isoluminance levels of all cell types when the threshold for excluding non-visual ROIs is varied, mean ±SEM across flies indicated by color line and shading, and black lines indicate the number of flies. Vertical gray lines indicate the thresholds used to exclude non-responding ROIs, and horizontal dashed lines indicate the corresponding mean ROI isoluminance levels calculated. **c**. FDR corrected *p*-values of two sample *t*-tests of isoluminance levels of pairs of cell types shown in Fig. 5f.

## Notes

### Competing Interest Statement

The authors have declared no competing interest.

